# Cadherin-13 is a critical regulator of GABAergic modulation in human stem cell derived neuronal networks

**DOI:** 10.1101/2020.05.07.082453

**Authors:** Britt Mossink, Jon-Ruben van Rhijn, Shan Wang, Eline J. H. van Hugte, Katrin Linda, Jitske Bak, Anouk H. A. Verboven, Martijn Selten, Alessia Anania, Sophie Jansen, Jason M. Keller, Teun Klein Gunnewiek, Chantal Schoenmaker, Astrid Oudakker, Monica Frega, Hans van Bokhoven, Dirk Schubert, Nael Nadif Kasri

## Abstract

Activity in the healthy brain relies on concerted interplay of excitation (E) and inhibition (I) via balanced synaptic communication between glutamatergic and GABAergic neurons. A growing number of studies imply that disruption of this E/I balance is a commonality in many brain disorders, however, obtaining mechanistic insight into these disruptions, with translational value for the human patient, has typically been hampered by methodological limitations. *Cadherin-13* (*CDH13*) has strongly been associated to attention-deficit/hyperactivity disorder and comorbid disorders such as autism and schizophrenia. CDH13 localises at inhibitory presynapses, specifically of parvalbumin (PV) and somatostatin (SST) expressing GABAergic neurons. However, the mechanism by which CDH13 regulates the function of inhibitory synapses in human neurons remains unknown. Starting from human induced pluripotent stem cells, we established a robust method to generate a homogenous population of SST and PV expressing GABAergic neurons (iGABA) *in vitro*, and co-cultured these with glutamatergic neurons at defined E/I ratios on micro-electrode arrays. We identified functional network parameters that are most reliably affected by GABAergic modulation as such, and through alterations of E/I balance by reduced expression of CDH13 in iGABAs. We found that CDH13-deficiency in iGABAs decreased E/I balance by means of increased inhibition. Moreover, CDH13 interacts with Integrin-β1 and Integrin-β3, which play opposite roles in the regulation of inhibitory synaptic strength via this interaction. Taken together, this model allows for standardized investigation of the E/I balance in a human neuronal background and can be deployed to dissect the cell-type specific contribution of disease genes to the E/I balance.

## Introduction

Neuronal network activity is controlled by a tightly regulated interplay between excitation (E) and inhibition (I). In the healthy brain, this interplay maintains a certain E/I ratio via balanced synaptic communication between glutamatergic and GABAergic neurons^1, 2^, resulting in the so called ‘E/I balance’. A growing number of studies imply that the E/I balance is disrupted in many neurodevelopmental disorders (NDDs)^3, 4^, including monogenic disorders, where the causative mutations are typically related to altered neuronal excitability and/or synaptic communication^5–7^, as well as polygenic disorders, such as autism spectrum disorders (ASD), attention deficit hyperactivity disorder (ADHD) and schizophrenia^4, 8^. Mutations in *Cadherin-13* (*CDH13*, also known as T-Cadherin or H-Cadherin)^9^ have been associated with ADHD^10–12^ and comorbid disorders such as ASD, schizophrenia, alcohol dependence and violent behaviour^13–16^. CDH13 is a special member of the cadherin superfamily since it lacks a transmembrane- and intracellular domain, and in contrast to other Cadherins, is anchored to the membrane via a glycosylphosphatidylinositol (GPI) anchor^17, 18^. Because of this relatively weak connection to the outer membrane^19^, CDH13 has been proposed to function as a regulatory protein, rather than an adhesion molecule^20, 21^. Indeed, CDH13 was shown to have a role in axon guidance and outgrowth^22–24^ as well as in regulation of apoptosis during cortical development^25^. CDH13 is expressed in different cell types, dependent on brain regions, including glutamatergic, GABAergic and serotonergic neurons^24, 26–28^. We recently showed that in the hippocampus CDH13 is located to the presynaptic compartment of inhibitory GABAergic neurons, specifically of parvalbumin (PV^+^) and somatostatin (SST^+^) expressing neurons, and that *Cdh13* knock-out mice (*Cdh13*^−/−^) show an increased inhibitory, but not excitatory synaptic input onto hippocampal CA1 pyramidal neurons^9^. In addition, these mice display deficits in learning and memory^9^. However, the mechanism via which CDH13 regulates GABAergic synapses remains unknown.

The E/I balance is particularly vulnerable to altered function and communication of GABAergic inhibitory neurons, whereas altered glutamatergic excitatory neuronal function often results in compensatory mechanisms that reinstate the E/I balance on the network level^2, 29^. Moreover, specific classes of GABAergic neurons, such as SST^+^ and PV^+^ neurons have been found to have a particularly strong influence on the E/I balance^30–32^. Although recent advances in differentiating human induced pluripotent stem cells (hiPSCs) into GABAergic neurons^33–42^, protocols that enable the generation of SST^+^ and PV^+^ human neurons are still challenging due to the long functional maturation of these cells^43^. Investigating E/I balance in human *in vitro* models for brain disorders ideally requires a model system that consists of a) neuronal networks with a known and reproducible composition of functional GABAergic and glutamatergic neuron classes, including PV^+^ and SST^+^ neurons, b) GABAergic signalling that matures to the functional state of shaping network behaviour by postsynaptic inhibition of neuronal activity, c) a neuronal network that allows controlling the ratio of glutamatergic and GABAergic neurons as well as cell-type specific manipulations of either cell type and d) the possibility to assess and manipulate the neuronal communication on single neuron as well as the larger scale neuronal network level.

In this study, we investigated the role of CDH13 in maintaining E/I balance in a human neuronal model. We describe a protocol that uses direct differentiation of hiPSCs into pure populations of either induced GABAergic or induced glutamatergic neurons through transcription factor-based reprogramming^36, 38, 39^. The induced GABAergic neurons included SST^+^ as well putative PV^+^ GABAergic neurons. When co-culturing these neurons with glutamatergic neurons over the course of seven weeks, they exerted inhibitory modulation of postsynaptic neurons, both on a single-cell and neuronal network level. We found that reducing *CDH13* expression specifically in human GABAergic neurons increases their inhibitory control onto human glutamatergic neurons. We further show that CDH13 is regulating GABAergic synapse function by heterophilic interactions with both integrin β1(ITGβ1) and integrin β3 (ITGβ3).

## Results

### Generation and characterization of human GABAergic neuron subtypes

We first developed a protocol for reproducibly generating and characterizing hiPSC-derived induced GABAergic neurons that can be co-cultured with induced glutamatergic neurons at predefined ratios. Specifically, we focused on generating SST^+^ and PV^+^ positive GABAergic neurons as CDH13 is highly expressed in these GABAergic subtypes. Moreover, these GABAergic subtypes are critical in the regulation of the E/I balance and have been implicated in NDDs^30–32^. By combining overexpression of *Ascl1*^38^ in hiPSCs paired with forskolin^36, 44^ (FSK, 10 μM) induction, we reliably generated GABAergic neurons (iGABA_A-FSK_, Figure 1a) that expressed the GABAergic neuronal markers glutamic acid decarboxylase 67 (GAD67) and γ-aminobutyric acid (GABA) at days in vitro (DIV) 49 (Figure 1b). When co-culturing iGABA_A-FSK_ neurons with iGLU_Ngn2_ neurons^39^ (Figure 1c), we identified an enrichment for SST (30%), calbindin CB (28%) and the PV-precursor protein MEF2C^45^ (17%) among the iGABA_A_-FSK neurons (Figure 1e). In addition, we found Synaptotagmin-2 (SYT2) positive puncta targeting the soma of glutamatergic neurons, indicative of PV^+^ synapses (Figure 1e, inset)^46^. Co-localization of the presynaptic vesicular GABA transporter (VGAT) and the postsynaptic scaffolding protein gephyrin indicated that inhibitory synapses are being formed on both the soma and dendrites (Figure 1d). RNAseq analysis at DIV 49 further confirmed that E/I networks highly express *SST*, *MEF2C* and genes expressed in mature fast spiking neurons (*FGF13*^47^, *LGL2*^47^, *PVALB*), as well as genes coding for Glutamate and GABA transporters (*SLC17A6/7*, *GAD1/2*) and GABAergic neuron development (*DLX1-6, LHX6, ZEB2, SOX6*, Figure 1f).

**Figure 1.**
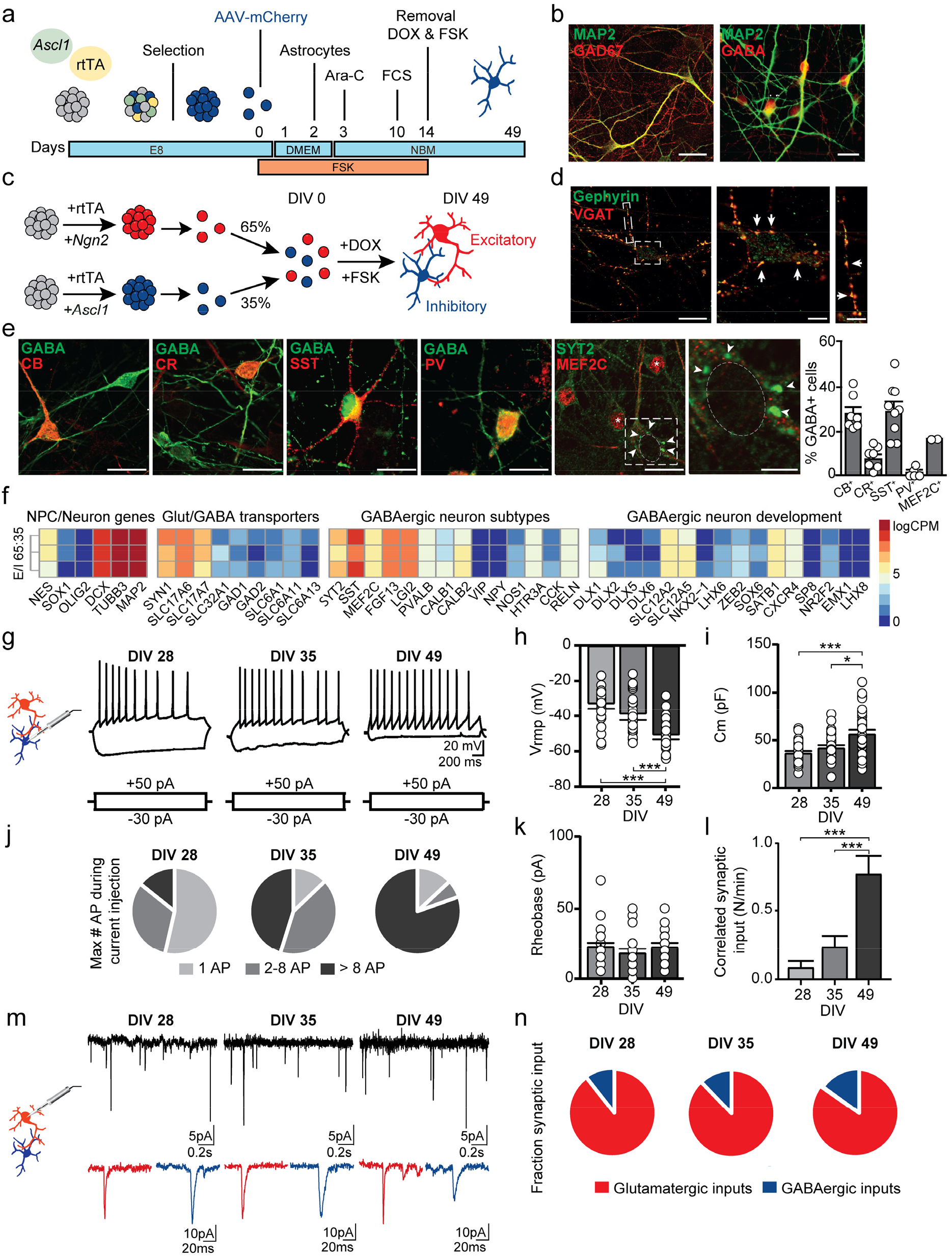
Rapid generation of human GABAergic neurons by overexpression of *Ascl1* and forskolin. (**a**) Culturing paradigm for the generation of induced GABAergic neurons (iGABAA-FSK). (**b**) iGABA_A-FSK_ neuron immunostaining at DIV 49 for neuronal marker MAP2 co-labelled with Glutamate decarboxylase (GAD) 67 or GABA. (**c**) iGABA_A-FSK_ are co-cultured from DIV 0 on with iGLU_Ngn2_ to promote functional maturation (named E/I networks), in a ratio of E/I 65:35. (**d**) VGAT and Gephyrin co-localisation in E/I networks at DIV 49. (**e**) Immunostaining for GABA co-labelled with either calbindin (CB), calretinin (CR), somatostatin (SST), parvalbumin (PV), MEF2C (asterix) or synaptotagmin-2 (SYT2, arrowheads) in E/I networks (Quantification sample size n= 7-9 coverslips per condition). (**f**) Heatmap showing expression of GABAergic/Glutamatergic transporters and – subtypes genes, and expression of genes important in GABAergic neuron development in E/I 65:35 networks at DIV 49 (3 biological replicates from one neuronal preparation). Data represents the log-transformed counts per million (logCPM). (**g**) Representative firing patterns of iGABA_A+FSK_ neurons at DIV 28, −35 and −49. (**h-i**) Analysis of iGABA_A+FSK_ membrane properties including (**h**) resting membrane potential (Vrmp) and (**i**) membrane capacitance (Cm). (**j-k**) Analysis of action potentials evoked by step-depolarization of iGABA_A-FSK_ membranes including (**j**) fractions of maximum number of action potentials, and (**k**) Rheobase. (**l**) Quantifications of correlated synaptic input (number of synaptic burst/minute). (**m**) Spontaneous glutamatergic (red inset) and GABAergic (blue inset) postsynaptic inputs (sPSCs) received by iGLU_Ngn2_. (**n**) Quantification of synaptic input types (Sample size for DIV 28 n=39, DIV 35 n=38, DIV 49 n=41 cells from 3 batches). All data represent means ± SEM. * p < 0.05; *** p < 0.001 (One-way ANOVA with Tukey correction for multiple testing was used to compare between DIVs). Scale bar is 20 μM, scale bars of zoom-in pictures are 6 μM.

Next, we functionally characterized the maturation of these composite E/I networks at DIV 28, 35 and 49. We visually identified iGABA_A-FSK_ neurons using mCherry labeling in single-cell patch-clamp recordings (Figure 1g-l, Supplementary figure 1a-f). At DIV 28 and later all recorded iGABA_A-FSK_ neurons could reliably elicit action potentials (APs, Figure 1g, j). As expected, during development we observed a hyperpolarization of the resting membrane potential (V_rmp_, Figure 1h), as well as an increase in membrane capacitance, indicating cell growth and maturation (Figure 1i). The rheobase remained unchanged (Figure 1k). No effect on the level of intrinsic properties was measured in iGLU_Ngn2_ neurons cultured in the presence of iGABA_A-FSK_ neurons in E/I networks (Supplementary figure 1g-p and q and Supplementary table 9).

In order to confirm that iGABA_A-FSK_ and iGLU_Ngn2_ functionally form an integrated network, we measured spontaneous GABAergic and glutamatergic synaptic inputs onto iGLU_Ngn2_ neurons (Figure 1l-n). By using decay time as a threshold to separate glutamatergic and GABAergic events^38, 48^ (see material and methods, Supplementary figure 1r-u and Supplementary table 1), we show that iGLU_Ngn2_ neurons received both spontaneous glutamatergic and GABAergic synaptic inputs (spontaneous postsynaptic currents, sPSC) throughout development when recorded at a membrane potential of −60 mV (i.e. at DIV 28, 35 and 49, Figure 1m). As a whole, during development we found a slight increase in the relative contribution of GABAergic inputs to all sPSCs (Figure 1n) and a significant increase in the number of spontaneous synchronized synaptic inputs (bursts) onto the iGLU_Ngn2_ neurons (Figure 1l), indicating robust integration of iGABA_A-FSK_ neurons into the E/I network as well as network-wide increased synaptic connectivity in the E/I networks over time^49^.

### Functional maturation of GABAergic synaptic responses in iGLU_Ngn2_ neurons

A hyperpolarizing shift in the chloride gradient dependent GABA reversal potential is key for enabling GABAergic synaptic inputs to modulate network activity by either shunting or hyperpolarizing inhibition and thus for establishing E/I balance during network development^50^. Local application of GABA onto iGLU_Ngn2_ somata during development revealed a prominent hyperpolarizing shift in the GABA reversal potential between DIV 35 and DIV 49 (Supplementary figure 2a-c). This hyperpolarizing shift of the GABA reversal potential has been shown in literature to be mediated through a decreased NKCC1:KCC2 chloride cotransporter expression ratio^50^. In accordance with literature, in our E/I networks the NKCC1:KCC2 expression ratio decreased between DIV 35 and 49 (Supplementary figure 2d-f and Supplementary table 2). Taken together, overexpression of *Ascl1* together with FSK supplementation leads to iGABA_A-FSK_ neuron induction enriched for SST^+^, CB^+^ and PV-precursor cell types, which by DIV 49 can exert a hyperpolarizing influence on iGLU_Ngn2_ neurons.

### iGABA_A-FSK_ show inhibitory control in E/I networks recorded by micro-electrode arrays

Having established a protocol for generating SST^+^ and PV^+^ iGABA_A-FSK_ neurons that can exert a hyperpolarizing (inhibitory) influence on iGLU_Ngn2_ neurons, we next investigated how these GABAergic neurons functionally modulate neuronal network development. We performed a comprehensive network analysis comparing two different network compositions of either iGLU_Ngn2_ alone (E/I ratio: 100:0), or in co-culture with iGABA_A-FSK_ neurons (E/I ratio: 65:35) on multi-electrode arrays (MEAs). Neuronal networks recorded on MEAs can display three distinctive patterns of activity, namely (i) random spiking activity (Figure 2a, green box), (ii) activity that is organized into a local burst (i.e. high frequency trains of spikes, Figure 2a, red box) and (iii) network wide bursting (i.e. bursts detected in all channels, Figure 2a, purple box) during development^51^. First, we confirmed that at DIV 49 treatment of E/I networks with 100 μM GABA completely abolished neuronal network activity (Supplementary figure 3a, b). Next, we compared the MEA recordings between the two network compositions side by side at DIV 35, 42 and 49 (Figure 2). Using discriminant analysis of 9 independent MEA parameters at all time points, we identified network burst duration (NBD) followed by network burst rate (NBR), mean firing rate (MFR) and the percentage of random spikes (PRS) as the main parameters that explain the significant differences in network activity between E/I 100:0 and E/I 65:35 networks (Figure 2d-f). Specifically, over development (i.e. at DIV 35, 42 and 49), in particular in E/I 65:35 we detected shortening of the NBD (Figure 2g), as well as a reduced NBR (Figure 2h) and MFR (Figure 2i), in contrast to an increased PRS (Figure 2j) as compared to E/I 100:0 networks. Interestingly, all of these network activity parameters only became significantly different between E/I 65:35 and E/I 100:0 after DIV 42 (Supplementary table 3). The time point for these differences to become significant indicates that the hyperpolarizing shift of the GABA reversal potential and thereby the maturation of the inhibitory system is underlying the different trajectories in functional network development between E/I 65:35 and E/I 100:0 networks. We found similar differences in these network parameters in E/I networks using an independent second *Ascl1* transduced control line (Supplementary figure 3c-d). Together, our results show that we can monitor and quantify the modulation of network activity by mature iGABA_A-FSK_ neurons during development on MEAs using a well-defined set of MEA parameters.

**Figure 2.**
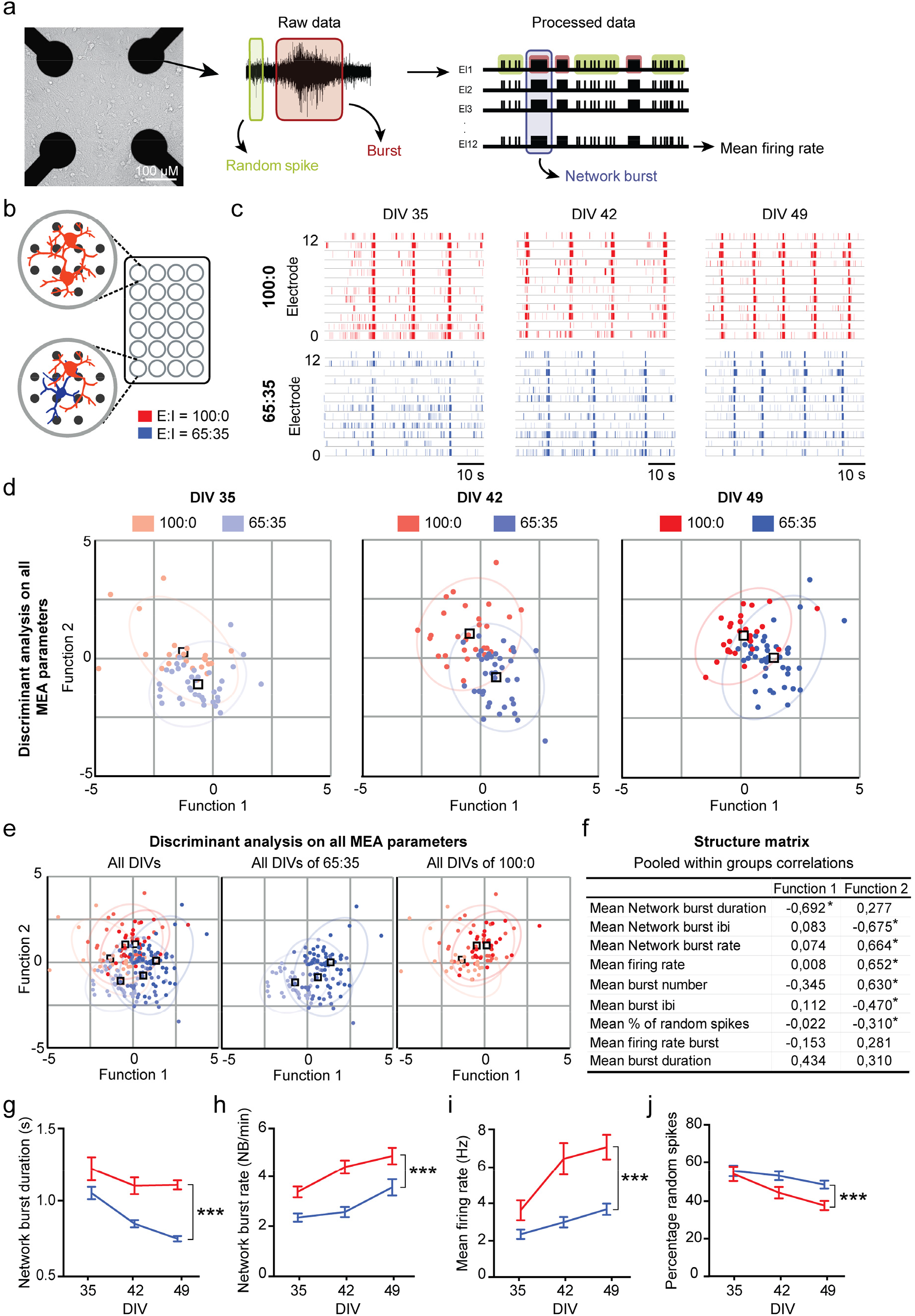
Discriminant analysis of E/I 100:0 and 65:35 networks reveal MEA parameters that reliably change depending on the hyperpolarizing GABA shift. (**a**) Representative image of neuronal density after plating E/I networks on micro-electrode arrays (MEAs). Schematic representation of the different types of spontaneous electric activity measured on MEAs. (**b**) iGLU_Ngn2_ alone (E/I ratio: 100:0) or in co-culture with iGABA_A-FSK_ (E/I ratio: 65:35) were recorded side by side on a multiwell MEA. (**c**) Representative raster plots showing 60 s of electrophysiological activity recorded from 12 electrodes of 100:0 (red) or 65:35 (dark blue) cultures at DIV 35, −42 and −49. (**d-e**) Canonical scores plots based on discriminant analyses of all 9 analysed MEA parameters for E/I 100:0 and 65:35 networks (**d**) at all DIVs separate, (**e**) all DIVs combined (left panel) only E:I 65:35 cultures at all DIV’s (second panel) and only E:I 100:0 cultures at all DIV’s (third panel). Discriminant functions are based on the following network activity parameters: firing rate, single channel burst rate, -duration, -firing rate in burst and -IBI, network burst rate, -duration and -IBI, percentage of random spikes. Ellipses are centred on the group centroids. (**f**) Structure matrix values showing which parameters explain the changes in neuronal network activity. Significantly changed parameters are marked with an Asterix. (**g-j**) Quantifications of neuronal network activity including (**g**) network burst duration, (**h**) network burst rate, (**i**) mean firing rate and (**j**) percentage of random spikes (Sample size n for 100:0 DIV 35 n= 25, DIV 42 n= 30, DIV 49 n= 29; 65:35 DIV 35 n= 40, DIV 42 n= 39 and DIV 49 n= 38 individual wells from 6 individual neuronal preparations). DIV: Days in vitro. All data represent means ± SEM. *** *p* < 0.001 (Mixed model Two-way ANOVA was performed between DIVs, p values were corrected for multiple comparisons using Sidak’s). IBI: Inter-burst interval.

### iGABA_A-FSK_ exhibit scalable inhibitory control onto the neuronal network

We evaluated to which extent the inhibition-mediated changes on the aforementioned MEA parameters depends on the specific ratio of iGLU_Ngn2_:iGABA_A-FSK_ present in our neuronal networks. To this end we co-cultured four different E/I ratios: 100:0, 95:5, 75:25 and 65:35 on MEAs and recorded spontaneous activity at DIV 49. In all conditions, the number of iGLU_Ngn2_ neurons was kept constant, whilst the number of iGABA_A-FSK_ was changed. Our data shows that the length of NBD was negatively correlated to the percentage of iGABA_A-FSK_ neurons (Figure 3a-e). Additionally, to the shortening of the NBD with increasing percentages of iGABA_A-FSK_ in the networks, we also detected NBDs to be composed of fewer detected spikes (Figure 3b-e). Furthermore, increasing percentages of iGABA_A-FSK_ in the networks led to a significant reduction in the MFR (Supplementary figure 3e) and NBR (Supplementary figure 3f) as well as an increase in PRS (Supplementary figure 3g and Supplementary table 4).

**Figure 3.**
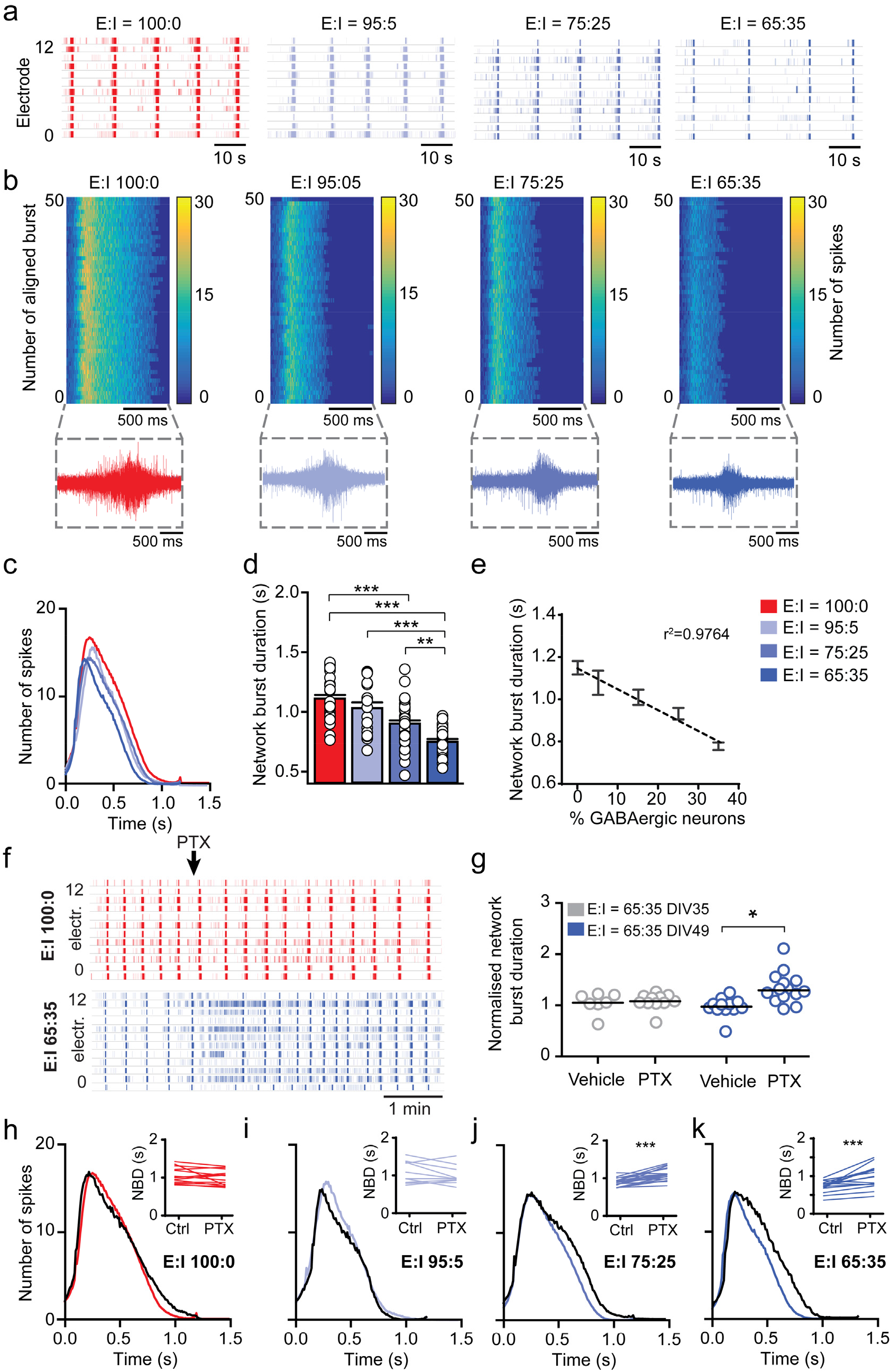
iGABA_A-FSK_ show scalable functional inhibition on the neural network at DIV 49. (**a**) Representative raster plots showing 60 s of electrophysiological activity recorded from E:I 100:0 (red), 95:5 (light blue), 75:25 (blue) or 65:35 (dark blue) networks at DIV 49. (**b**) Representative network burst alignment from one recording of E:I 100:0, 95:5, 75:25 or 65:35 networks, colour code represents the spikes. Inset: representative network burst. (**c**) Average network burst shape of representative cultures from 100:0, 95:5, 75:25 or 65:35 networks at DIV 49 (Sample size n for 100:0 cultures n=20, 95:5 n=12, 75:25 n=23 and 65:35 n=26 individual wells. For E:I 65:35 networks *p*=0.008, multiple t-test on bins using Holm-Sidak method). (**d**) Quantification of the average network burst duration of E:I 100:0, 95:5, 75:25 and 65:35 networks (Sample size for 100:0 n=29, 95:5 n=20, 75:25 n=38 and 65:35 n=38 individual wells, Kruskal-Wallis Two way ANOVA was performed between ratio’s at DIV 49, corrected for multiple testing using Dunn’s method). (**e**) Linear regression plot of the average network burst duration from 100:0, 95:5, 85:15, 75:25 or 65:35 cultures at DIV 49 (y=−9.628x+1109, *p*=0.0119). (**f**) Representative raster plots of 5 minutes showing the effect of acute 100 μM picrotoxin (PTX) treatment on E/I 100:0 and 65:35 networks at DIV 49. (**g**) Normalised network burst duration of E/I 65:35 networks treated acutely with vehicle or PTX at DIV 35 and DIV 49, normalised to their respective baseline recording (Sample size n for DIV 35 + vehicle n= 8; DIV 35 + PTX n=11; DIV 49 + vehicle n=12 and DIV 49 + PTX n=15 individual wells, Mann-Whitney test with post hoc Bonferroni correction for multiple testing was performed). (**h-k**) Quantification of network burst shape after acute PTX treatment in (**h**) 100:0, (**i**) 95:5, (**j**) 75:25 and (**k**) 65:35 cultures at DIV 49 (black line indicates the average burst shape of wells treated with PTX, sample size n for 100:0 cultures n=9, 95:5 n=6, 75:25 n=12 and 65:35 n=11 individual wells, 100:0 *p*=0.5582, 95:5 *p*= 0.1857, 75:25 *p*=0.1050 and 65:35 *p*= 0.0013, multiple t-test on bins using Holm-Sidak method). Inset: Paired t-test of the mean network burst duration before and after treatment with PTX (Sample size n for 100:0 cultures n=15, 95:5 n=10, 75:25 n=19 and 65:35 n=15 individual wells). DIV: Days in vitro. All data represent means ± SEM. * p < 0.05, * p < 0.01, *** p < 0.001.

We showed that iGABA_A-FSK_ neurons shape network burst activity at DIV 49 through inhibition. We next investigated how acute removal of inhibitory control alters the NBD compared to networks that lacked inhibitory control during development (i.e. iGLU_Ngn2_ only cultures). Following acute treatment with either vehicle or 100 μM Picrotoxin (PTX), the NBD did not change in the 100:0 E/I networks at DIV 49 (Figure 3f, top panel). In contrast, PTX significantly increased the NBD and MFR of 65:35 E/I networks (Figure 3f, bottom panel). In accordance with our data on the hyperpolarizing shift of the GABA reversal potential during development, in these 65:35 E/I networks acute treatment with PTX at DIV 35 did neither affect the NBD (Figure 3g) nor the MFR (Supplementary figure 3h). Moreover, the impact of PTX treatment on the NBD and MFR was again scalable to the ratio of iGABA_A-FSK_ neurons present (Figure 3h-k and Supplementary figure 3i). Finally, we infected E/I networks with an AAV expressing Channelrhodopsin-2 in either iGLU_Ngn2_ or iGABA_A-FSK_ neurons. Optogenetic activation of iGLU_Ngn_ neurons at DIV 49 resulted in an increase in MFR (Supplementary figure 3j), whereas optogenetic activation of iGABA_A-FSK_ neurons reduced the MFR (Supplementary figure 3k and Supplementary table 5). Together, these data show that iGABA_A-FSK_ neurons at the network level exert robust inhibitory control at DIV 49.

### Knockdown of *CDH13* increases inhibitory control onto neuronal networks

To investigate the role of *CDH13* in maintaining E/I balance in human neurons, we first verified its expression in 65:35 E/I networks. We found that amongst many other disorder-related genes with a prominent influence on the E/I balance such as Neuroligin (*NLGN*) and Neurexin (*NRXN*)^52^, also *CDH13* to be expressed in these E/I networks (Supplementary figure 4a). Moreover, we found CDH13 to be co-localized with VGAT and SYT2 (Figure 4a), demonstrating that, as in rodent neurons^9^, also in human iPSC derived E/I networks CDH13 is localized to inhibitory presynapses. Of note, iGLU_Ngn2_ only networks did not express *CDH13*, confirming that *CDH13* is exclusively expressed in iGABA_A-FSK_ neurons (Supplementary figure 4b).

**Figure 4.**
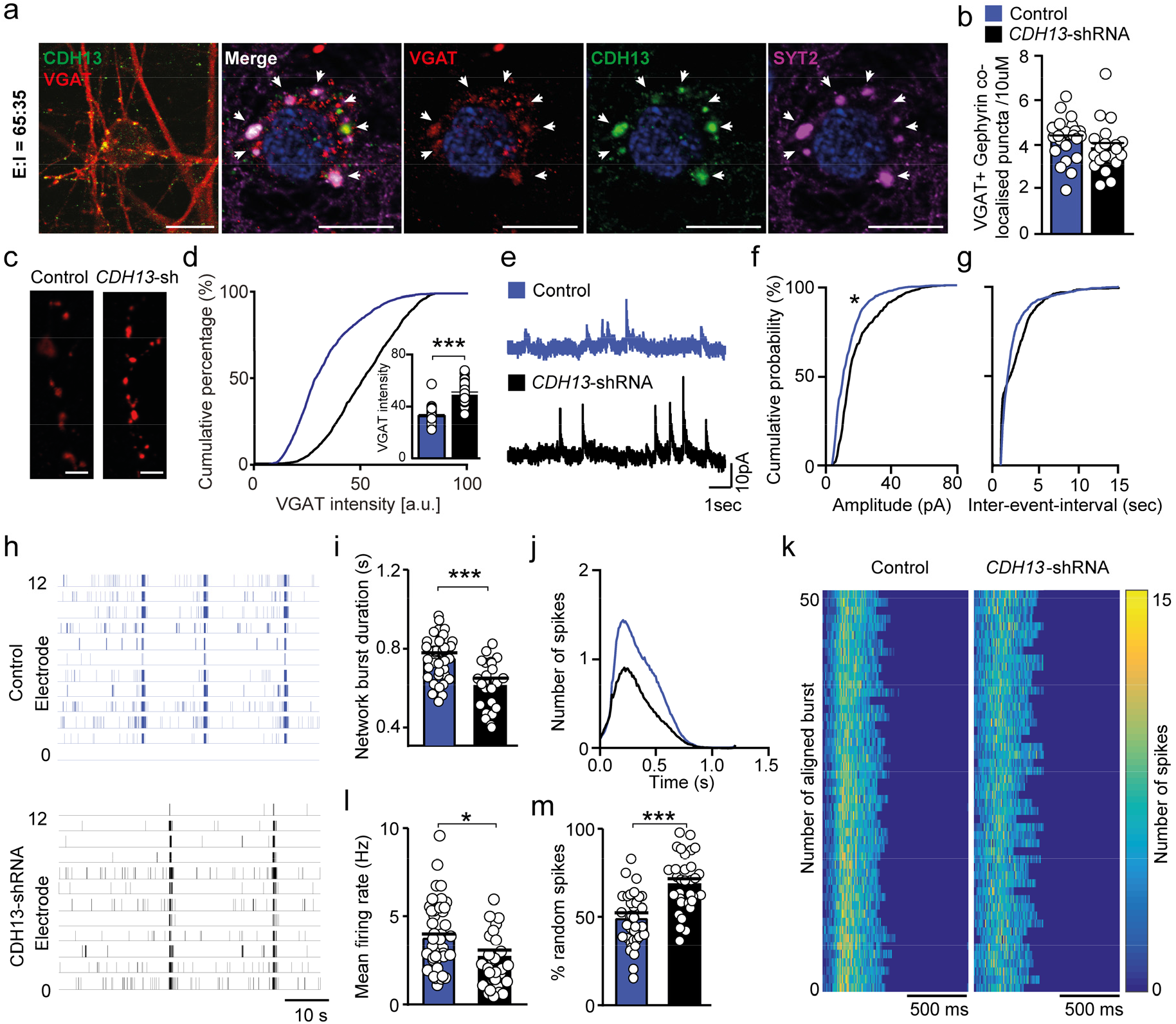
Knockdown of CDH13 in iGABA_A-FSK_ leads to increased inhibition in E/I networks. Co-labelling of VGAT (red), CDH13 (green) and SYT2 (purple) in E/I 65:35 controls at the inhibitory presynapse (scale bar 10 μM). (**b**) Number of VGAT and Gephyrin co-localised puncta in E/I 65:35 control and CDH13-deficient networks (control n=23, CDH13-deficient n=21 analysed cells from 3 individual neuronal preparations). (**c**) Representative VGAT staining in E/I 65:35 control and CDH13-deficient networks at DIV 49 (scale bar 6 μM). (**d**) Quantification of VGAT intensity (arbitrary units) per puncta (control n=25 and CDH13-deficient n= 26 cells from 3 individual neuronal preparations). (**e**) Example trace of sIPSC activity in E/I 65:35 control and CDH13-deficient networks at DIV 70. (**f**) Cumulative distribution of sIPSC amplitude and (**g**) frequency in E/I 65:35 control and CDH13-deficient networks (control n=7, CDH13-deficient n=10 recorded cells from 2 individual neuronal preparations). (**h**) Representative raster plots showing 60 s of electrophysiological activity recorded from E/I 65:35 control and CDH13-shRNA treated cultures at DIV 49. (**i**) Average network burst duration in E/I 65:35 control and CDH13-deficient networks. (**j**) Average network burst shape of representative cultures from E/I 65:35 control and CDH13-deficient networks at DIV 49 (*p*=0.00071, Multiple t-test on bins using Holm-Sidak method, Sample size for control n=26 and CDH13-deficient n=12 individual wells). (**k**) Representative network burst alignment from one recording of E/I 65:35 control and CDH13-deficient networks, colour code represents the number of spikes. (**l**) Mean firing rate and (**m**) percentage of random spikes in E/I 65:35 control and CDH13-deficient networks (Panel i, l and m sample size n for control n=38 and 65:35 CDH13-deficient wells n=33 individual wells from 3 neuronal preparations. Mann-Whitney test with post hoc Bonferroni correction for multiple testing was performed). All data represent means ± SEM. * p < 0.05; *** p < 0.001. DIV: Days in vitro.

After confirming *CDH13* expression in control 65:35 E/I networks, we investigated the functional consequences of reduced *CDH13* expression. To this end we employed validated short hairpin RNAs (shRNA) to downregulate *CDH13* expression^28^, specifically in iGABAA-FSK neurons by only infecting *Ascl1* expressing hiPSCs prior to co-culturing (Supplementary figure 4c and Supplementary table 10). At DIV 49 CDH13-deficient networks showed neither changes in the number of inhibitory presynapses identified by VGAT labelling (Supplementary figure 4d), nor in inhibitory synapses identified by juxtaposed VGAT/Gephyrin puncta (Figure 4b), when comparing to control networks treated with scrambled hairpin. However, the CDH13-deficient networks showed a striking increase in the mean intensity of VGAT puncta (Figure 4c, d), suggesting that loss of CDH13 did not affect the number of synapses, but rather increased inhibitory strength. We confirmed this by measuring GABAergic sPSCs in iGLU_Ngn2_ cells, where we found increased GABAergic sPSC amplitudes but no changes in sPSC frequency in CDH13-deficient networks (Figure 4e-g and Supplementary table 6), further supporting that CDH13 is a negative regulator of inhibitory synaptic function.

Next, we assessed the impact of CDH13-deficiency in iGABA_A-FSK_ neurons on the level of network activity at DIV 49 (Figure 4h). Lentiviral infection as such did not affect network activity of control E/I networks (non-treated vs scrambled hairpin, Supplementary figure 4f). We found reduced NBD (Figure 4i) together with altered average burst shape and less detected spikes within a network burst (Figure 4j, k) in CDH13-deficient networks. Furthermore, the CDH13-deficient networks showed a significantly reduced MFR (Figure 4l), while the PRS was significantly increased (Figure 4m and Supplementary table 6). We also found a trend towards a lower network burst rate in CDH13-deficient networks as compared to controls at DIV 49 (Supplementary figure 4e, *p*=0.0649). All these changes in network parameters are consistent with an increase in inhibition upon loss-of-function of CDH13.

### CDH13 regulates inhibitory synaptic strength via interaction with ITGβ1 and ITGβ3

The observed increase of VGAT expression in CDH13-deficient networks implies that CDH13 is a negative regulator of synapse function, however, the underlying mechanism is unknown. CDH13 is a GPI-anchored protein, which suggests that binding to other membrane bound proteins is required to exert its function^15^. In agreement with rodent data^9^, we showed that in human iPSC derived E/I networks CDH13 expression is restricted to GABAergic neurons; therefore a heterophilic interaction is likely to be required for CDH13 to exert its function. Previous co-immunoprecipitation studies in endothelial cells identified the GABA-A receptor α1 subunit (GABAAα1) and ITGβ3^53^ as potential interaction partner for CDH13. Overexpression of CDH13 has also been shown to increase ITGβ1 expression in squamous carcinoma cells^54^, even though a direct interaction has not been reported. Interestingly, ITGβ1 and ITGβ3 have opposite functions in regulating synaptic dwell time of glycine receptors in spinal cord neurons, bidirectionally regulating the synaptic strength of these inhibitory synapses^55^. Both ITGβ1 and ITGβ3 are expressed in glutamatergic neurons, where they play a role in regulating glutamatergic synaptic function though the modulation of AMPARs^56–58^. However, until now a role in the regulation of GABAergic synaptic function in glutamatergic neurons has not been described for these integrins. PV^+^ synapses are enriched for the GABA receptor subunit α1 (GABAAα1^59^). Therefore, we hypothesised that CDH13 may play a role in regulation of GABAergic synapse stability via direct interaction with GABAAα1, ITGβ1 or ITGβ3. We first assessed the cellular localisation of CDH13, ITGβ1, ITGβ3 and GABAAα1 in our E/I networks (Figure 5a-c). Whereas CDH13 co-localised with VGAT (Figure 4a) in the presynaptic terminal, GABAAα1 localised juxtapose of CDH13 (Figure 5a) and ITGβ1 (Figure 5b) and ITGβ3 (Figure 5c) localised juxtapose of VGAT, suggestive of a postsynaptic localisation. Next, we confirmed the interactions between ITGβ1, ITGβ3, GABAAα1 and CDH13 by co-immunoprecipitation experiments, using lysates from a human embryonic kidney cell line (HEK293) expressing GFP-tagged GABAAα1, GFP-tagged ITG β1 or ITGβ3 and myc-tagged CDH13 (Figure 5d-f).

**Figure 5.**
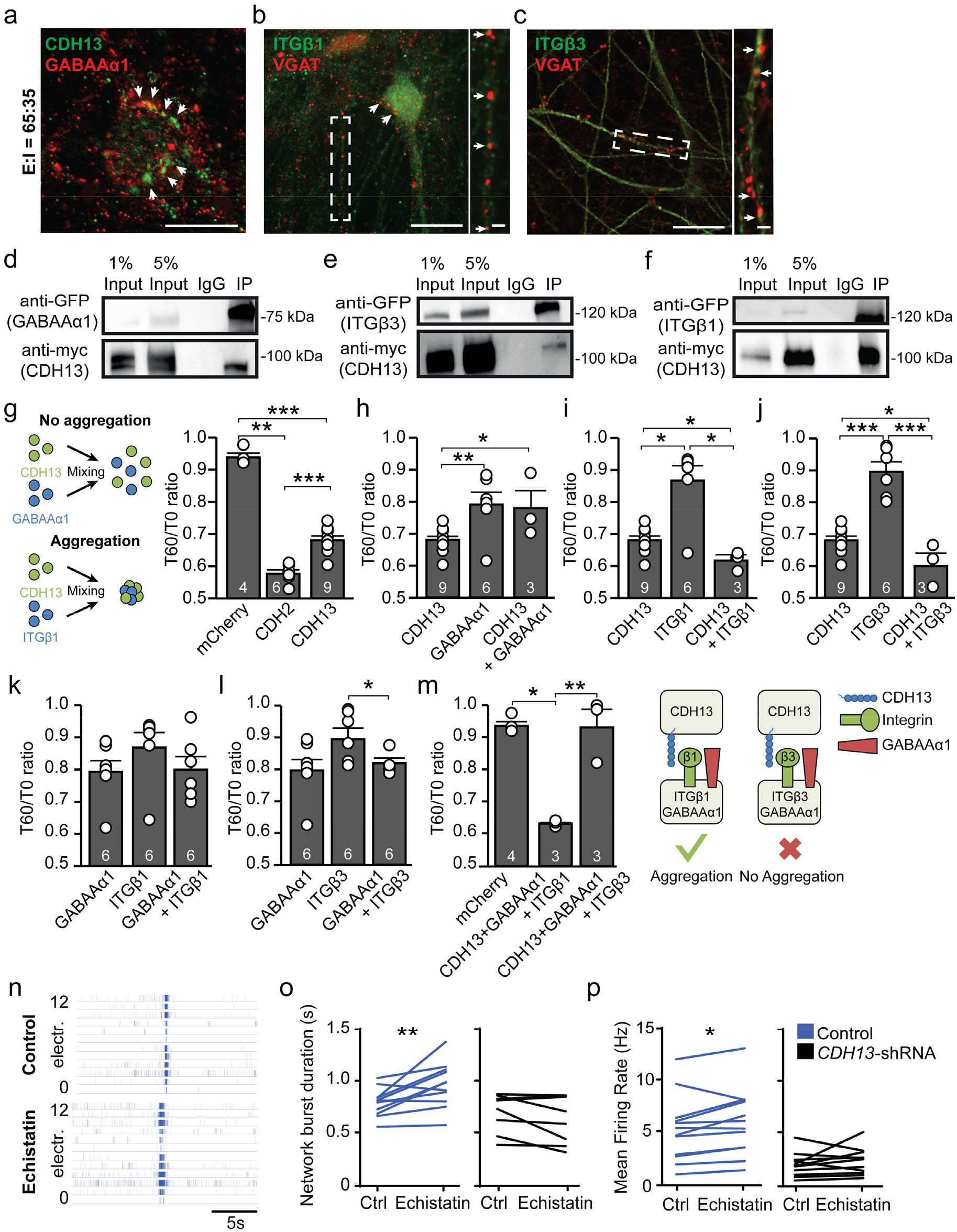
CDH13 interacts with integrin β1 (ITGβ1) and ITGβ3 *in vitro*. (**a-c**) Representative colocalisations of (**a**) GABAAα1 with CDH13, and (**b**) Integrin (ITG) β1 and (**c**) ITGβ3 with VGAT in E:I 65:35 networks. Scale bar is 20 μM, scale bars of zoom-in pictures are 6 μM. (**d-f**) Western blot showing coimmunoprecipitation of (**d**) CDH13 with GABAAα1, (**e**) CDH13 with ITGβ3 and (**f**) CDH13 with ITGβ1 in HEK cells. (**g-m**) Quantification of cell aggregation for indicated proteins in HEK cells (Sample size n represented in figure). (**n**) Representative rasterplots of E:I 65:35 control and CDH13-deficient networks treated with 100 μM Echistatin. (**o-p**) Quantification of (**o**) network burst duration and (**p**) mean firing rate of pre- and post echistatin treated E:I 65:35 control and CDH13-deficient networks (Sample size n for control n=12 and 65:35 CDH13-deficient wells n=15 individual wells from 3 neuronal preparations. Paired T-test with was performed between pre and post echistatin treatment conditions). All data represent means ± SEM. * p < 0.05; ** p < 0.01; *** p < 0.001. IP: Immunoprecipitation.

### Differential roles for ITGβ1 and ITGβ3 in cell adhesion assays

Since our data indicate that CDH13, ITGβ3 and GABAAα1 co-localise at the same synapse, we wanted to know if these proteins are able to play a role in cell adhesion. To this end we used a cell adhesion assay^60^. In this assay, we transfected HEK293T cells with a vector expressing CDH13, ITGβ1, ITGβ3 or GABAAα1 and quantified the degree of aggregation at two different time points and calculated the ratio (T60/T0 ratio). As negative and positive control for cell adhesion we transfected HEK293T cells respectively with mCherry or Cadherin 2 (CDH2), of which the relative strengths of binding are known (Figure 5g)^19, 61^. In line with literature, CDH2 showed a strong aggregation^61, 62^ (Figure 5g, low T60/T0: 0.58 ± 0.01), whereas mCherry-expressing HEK293T cells showed very little aggregation (Figure 5g, high T60/T0: 0.94 ± 0.01). Homophilic interactions of CDH13 have been proposed, but are predicted to be weak compared to CDH2 homophilic interactions^17, 19^. Indeed, in our assay CDH13 expressing HEK293T cells showed an intermediate value (Figure 5g and Supplementary table 7) and show that this assay has the sensitivity to distinguish between different strengths of cell-adhesion.

We then investigated the interactions between CDH13, ITGβ1, ITGβ3 or GABAAα1. GABAAα1 was co-transfected with GABAAβ3 to ensure surface expression of these proteins^63^. GABAAα1/β3 expressing HEK293T cells showed a weak homophilic interaction (Figure 5h). We then combined HEK293T cells expressing CDH13 or GABAAα1/β3, however the resulting T60/T0 ratio ni dicated no heterophilic adhesion between these proteins (Figure 5h and Supplementary table 7). Conversely, while ITGβ1 shows no homophilic interaction, consistent with previous reports^64^, CDH13 and ITGβ1 showed heterophilic interaction (Figure 5i). ITGβ3 showed the same pattern as ITGβ1, displaying a stronger interaction with CDH13 (Figure 5j) than either CDH13 or ITGβ3 alone. We next assessed the interaction between integrins and GABAα1/β3 in transsynaptic conformation. A mix of ITGβ1 expressing and GABAα1/β3 expressing cells did not show interaction (Figure 5k). We also found no interaction between ITGβ3 and GABAAα1/β3 (Figure 5l).

Finally, we investigated the aggregation using a protein arrangement as expected *in vivo*. We expressed either ITGβ1 or ITGβ3 together with GABAAα1/β3 in one population of HEK293T cells, representing the postsynaptic side. To represent the presynaptic side, we transfected HEK293T cells with CDH13 (Figure 5m). Surprisingly, we found that while ITGβ1/ GABAAα1/β3 expressing cells displayed a strong interaction with CDH13 expressing cells, ITGβ3/ GABAAα1/β3 expressing cells did not interact with CDH13 expressing cells (Figure 5m, Supplementary table 7). In conclusion, while both ITGβ1 and ITGβ3 show interaction with CDH13, co-expression of GABAAα1/β3 with the integrins leads to a loss of interaction between CDH13 and ITGβ3, specifically.

If ITGβ1 and ITGβ3 play a role in inhibitory synapse stabilization via their interaction with CDH13, disruption of integrin function should affect inhibitory transmission. In order to test this on the functional level we applied 100 nm Echistatin, an inhibitor of ITGβ1 and ITGβ3^65^, to E/I networks recorded at DIV 49 on MEA. Blocking ITGβ1/3 interaction increased NBD and MFR in control E/I networks, indicating that ITGβ1/3 play a role in maintaining inhibitory strength in control networks. Interestingly, Echistatin had no effect on CDH13-deficient networks compared to vehicle treated cells (Figure 5n-p and Supplementary table 8). Together, these data indicate that ITGβ1/ITGβ3 play a critical role in inhibitory synapse maintenance, and that this role is dependent on the presence of CDH13.

## Discussion

In this study, we describe a human *in vitro* neuronal model system for investigating the function of CDH13 in the maintenance of E/I balance. Loss of CDH13 in humans is linked to the aetiology of several NDDs, including ADHD and ASD^27, 66^, and in rodents has been found to alter E/I balance on the single-cell level^9^. Since the first postulation of an increased E/I ratio in ASD^67^, an increasing amount of studies has shown that altered E/I balance contributes to many NDDs^1^. Interestingly, evidence from both animal models and human studies suggest that altered function of PV^+^ GABAergic neurons is a common unifying pathway for common forms of NDDs^4, 30, 43^. Although several efforts have been made to generate PV^+^ GABAergic neurons from hiPSCs, their generation has been proven challenging^43^. Here we show that *Ascl1* overexpression and FSK supplementation resulted in approximately 30% SST^+^ GABAergic neurons. Even though a large population of distinct PV expressing neurons was absent, 15 to 20% of the GABAergic neurons were expressing MEF2C, a marker for immature PV^+^ neurons^45^. Together with the existence of soma targeting SYT2-positive GABAergic synapses onto iGLU_Ngn2_ neurons in our cultures, this implies that these MEF2C^+^ neurons represent PV^+^ precursor cells^46^.

After establishing a protocol that generates a reproducible composition of GABAergic neuronal classes, we confirmed the functional maturation of GABA signalling in the E/I networks. *In vivo*, the emergence of functional GABAergic inhibition via GABAA receptors is facilitated by a hyperpolarizing shift in the chloride reversal potential during development mediated through activity-dependent increase in the ratio of KCC2:NKCC1 chloride co-transporter expression in neurons^68^. Multiple studies have evaluated the generation of iGABA neurons based on the expression of GABAergic markers and synaptic GABA release^33–38, 40^. However, to our knowledge it has not been shown before that using direct differentiation of hiPSC into composite E/I networks, iGABA_A-FSK_ develop into neurons that functionally modulate iGLU_Ngn2_ network activity by GABA mediated postsynaptic shunting inhibition and/or hyperpolarizing inhibition. This is not only important for network phenotyping, but is also essential for iGLU_Ngn2_ maturation and the maintenance of the E/I balance^69–71^. Our data demonstrate that the generated E/I networks receive glutamatergic as well as GABAergic synaptic inputs and indeed show an decrease in the NKCC1:KCC2 ratio during development. At the functional level we could correlate this with a hyperpolarizing shift of the GABA reversal potential, indicating iGABA_A-FSK_ neurons in mature *in vitro* E/I networks can functionally modulate network activity in E/I networks.

This leaves the question regarding how to assess E/I balance at a neuronal network level. One well established model to generally assess neuronal network activity *in vitro* are cultures growing on MEAs^72–77^. Indeed, MEAs have shown to be a powerful tool to elucidate the contribution of receptors of excitatory and inhibitory synaptic transmission to spontaneous network activity in rodent *in vitro* cultures^78^. Here we show the development of human iPSC derived E/I networks over time, and describe network parameters that illustrate the modulation of hyperpolarizing/shunting inhibition by iGABA_A-FSK_ neurons. In relation to the temporal aspects of the hyperpolarizing shift in the chloride-gradient dependent GABA reversal potential, we show a decrease of the NBD, MFR and NBR and an increase in the PRS over development from DIV 35 to 49, which are in line with previously published work in rodent and human E/I networks on MEA^72, 76^. In particular, the shortening of the NBD has been demonstrated to be a hallmark of mature GABA mediated signaling in neuronal networks^76, 78, 79^, mainly by reducing the intra burst activity which in turn scales down the Mg^2+^ block release from the NMDAR pore^78, 80^. In our E/I cultures, we could not only reproduce the maturation dependent effects of GABAergic signaling on network bursts, but also demonstrated that these effects are scalable to the amount of inhibitory neurons in the E/I cultures: we were able to show a direct correlation between the different network parameters and the amount of inhibition.

Using this E/I network model, we studied the cell-type specific contribution of CDH13 in iGABA_A-FSK_ neurons. When comparing control networks with networks in which CDH13 expression is specifically reduced in only iGABA_A-FSK_ neurons, we found that CDH13-deficiency increased inhibitory control at the network level, which is in line with the synaptic phenotypes found in hippocampal CA1 neurons of *Cdh13*^−/−^ mice^9^. With keeping the scalable consequences of the amount of GABAergic neurons on network behaviour in mind, E/I cultures with CDH13 deficient iGABA_A-FSK_ neurons clearly imply an elevated impact of GABAergic signalling on the E/I cultures. One prominent feature illustrating the elevated impact of GABAergic signalling was the strong shortening of NBD, most likely mediated by elevated suppression of within burst spiking and consequently the suppression of late NMDAR dependent phase of the bursts^80^.

At the molecular level, we show that CDH13 co-immunoprecipitates with ITGβ1 and ITGβ3, and that CDH13 has the ability to bind both ITGβ1 and ITGβ3 in the cell adhesion assay. Interestingly, while co-expression of GABAAα1/β3 does not affect the interaction between CDH13 and ITGβ1, coexpression of GABAAα1/β3 with ITGβ3 completely abolished the interaction between CDH13 and ITGβ3. Both ITGβ1 and ITGβ3 are expressed by pyramidal neurons^56, 57^, and we show that these are expressed postsynaptically together with GABAAα1. This points to the intriguing possibility that ITGβ1 and ITGβ3 could function as a molecular switch for synapse maintenance. A similar function for ITGβ1/ITGβ3 has already been described previously in spinal cord neurons, where these integrins have opposite functions in the regulation of synaptic dwell time of glycine receptors through stabilisation (ITGβ1) and destabilization (ITGβ3) of the inhibitory synaptic scaffold protein gephyrin^55^, and via this mechanism regulate the strength of glycinergic synapses. The differential function of ITGβ1/ITGβ3 would allow glutamatergic neurons to control the amount of inhibitory input they receive. Since both ITGβ1 and ITGβ3 are also expressed in glutamatergic synapses, ITGβ1/ITGβ3 might be in the ideal position to maintain the E/I balance by regulating simultaneously the E and I input, respectively by stabilizing the excitatory and inhibitory postynaptic receptors^56, 57, 81^. It has recently been shown that cortical pyramidal neurons receive a relative amount of inhibitory synaptic input from GABAergic PV^+^ neurons corresponding to the excitatory drive onto that pyramidal neuron, thereby maintaining their E/I balance^82^. Since individual GABAergic PV^+^ neurons can differentially regulate their inhibitory strength onto individual postsynaptic pyramidal neurons^82^, it is likely that pyramidal neurons instruct the regulation of inhibitory synapses onto themselves. The complex of CDH13, ITGβ1 and ITGβ3 could play a role in this regulation. Loss of CDH13 would lead to the inability of the postsynaptic glutamatergic neuron to regulate inhibitory synapses formed onto itself via regulation of the ITGβ1/ITGβ3 ratio. Indeed, in *Cdh13*^−/−^ mice we previously reported an increase in inhibitory synapses^9^. The importance of CDH13 in this mechanism is underlined by our finding that while Echistatin affected neuronal network activity of control networks, it has no effect in CDH13-deficient networks. Future experiments should dissect the precise contribution of these proteins in synaptic regulation and stability.

## Methods

### Neuronal differentiation

HiPSCs from control #1 and #2 were differentiated into Glutamatergic cortical Layer 2/3 neurons by overexpressing mouse neuronal determinant Neurogenin 2 (*Ngn2*) upon doxycycline treatment^39, 77^ (referred to as *Ngn2* #1 and *Ngn2* #2, respectively). If not mentioned differently, all experiments were performed using the *Ngn2* #1 background.

GABAergic neurons were derived from control hiPSC line #2 and #3 by overexpressing mouse neuronal determinant Achaete-scute homolog 1 (*Ascl1*, plasmid was custom designed and cloned by VectorBuilder and is available upon request) upon doxycycline treatment with supplementation of Forskolin (10 μM, Sigma) (referred to as *Ascl1* #1 and *Ascl1* #2, respectively). If not mentioned differently, all experiments were performed using the *Ascl1* #1 background throughout this study. Glutamatergic neurons were either cultured alone or in co-culture with iGABA_A-FSK_. When co-cultured, GABAergic neurons were plated at days in vitro (DIV) 0 and labelled with AAV2-hSyn-mCherry (UNC Vector Core) for visualisation, with AAV2-hSyn-hChR2(H134R)-mCherry (UNC Vector Core) for optogenetic activation, or with lentivirus expressing GFP scrambled short hairpin RNA (shRNA, control) or *CHD13*-shRNA. After 5 hours of incubation, cultures were washed twice with DMEM/F12 (Thermo Fisher Scientific) before iGLU_Ngn2_ were plated on top. When changing the E/I ratio from 95:5, 85:15, 75:25 to 65:35, the number of iGLU_Ngn2_ present in the culture was always kept at a similar density whereas the number of iGABA_A-FSK_ was increased to make sure baseline electrophysiological activity was kept constant. HiPSCs were plated in E8 flex supplemented with doxycycline (4 μg/ml), revitacell (1:100, Thermo Fisher Scientific) and Forskolin. At DIV 1 cultures were switched to DMEM/F12 containing Forskolin (10 μM, Sigma), N2 (1:100, Thermo Fisher Scientific), non-essential amino acids (1:100, Sigma), primocin (0.1 μg/ml), NT3 (10 ng/ml), BDNF (10 ng/ml), and doxycycline (4μg/ml). To support neuronal maturation, freshly prepared rat astrocytes^77^ were added to the culture in a 1:1 ratio at DIV 2. At DIV 3 the medium was changed to Neurobasal medium (Thermo Fisher Scientific) supplemented with Forskolin (10 μM, Sigma), B-27 (Thermo Fisher Scientific), glutaMAX (Thermo Fisher Scientific), primocin (0.1 μg/ml), NT3 (10 ng/ml), BDNF (10 ng/ml), and doxycycline (4 μg/ml). Moreover, cytosine-b-D-arabinofuranoside (Ara-C; 2 μM; Sigma) was added once to remove any proliferating cell from the culture. From DIV 6 onwards half of the medium was refreshed three times a week. The medium was additionally supplemented with 2,5% FBS (Sigma) to support astrocyte viability from DIV 10 onwards. After DIV 13, Forskolin and doxycycline were removed from the culture medium. Neuronal cultures were kept through the whole differentiation process at 37°C/ 5% CO_2_. We compared the generation of E/I networks of 2 *Ascl1* stable control lines (i.e. composed of *Ngn2* #1 + *Ascl1* #1 or *Ngn2* #1 + *Ascl1* #2). The network activity on MEA (Supplementary figure 3e, f), single-cell recordings and immunohistochemistry analysis were not affected by combining different *Ascl1* stable lines. Therefore, all data gathered from *Ngn2* #1 + *Ascl1* #1 or *Ngn2* #1 + *Ascl1* #2 as mentioned above were pooled in the respective analysis.

### RNA interference

For RNAi knockdown experiments, DNA fragments encoding shRNAs directed against human *CDH13* mRNA (Sigma) were cloned into the pTRIPΔU3-EF1α-EGFP lentiviral vector. The hairpin sequences are listed in Supplementary table 12. Empty vector expressing GFP only was used as control vector. Lentiviral particles were prepared, and tittered as described previously^83^. In brief, lentiviruses were generated by co-transfecting the transfer vector, the psPAX2 packaging vector (Addgene #12260), and the VSVG envelope glycoprotein vector pMD2-G (Addgene plasmid #12259) into HEK293T cells, using calcium phosphate precipitation. Supernatants of culture media were collected 48 hours after transfection and filtered through a 0.45 μm syringe filter. Viral particles were then stored at −80°C until use.

### Micro-electrode array recordings and data analysis

All recordings were performed using the 24-well MEA system (Multichannel Systems, MCS GmbH, Reutlingen, Germany) as described before^75, 77^. Spontaneous electrophysiological activity of E/I networks was recorded for 10 min at 37°C and constant flow of humidified gas (5% CO_2_ and 95% O_2_). The raw signal was sampled at 10 kHz and filtered with a high-pass filter (i.e. 2^nd^ order Butterworth, 100 Hz cut-off frequency) and a low-pass filter (i.e. 4^th^ order Butterworth, 3500 Hz cut-off frequency). The threshold for detecting spikes was set at ± 4.5 standard deviations. We performed off-line data analysis by using Multiwell Analyzer (i.e. software from the 24-well MEA system that allows the extraction of the spike trains) and in-house algorithms in MATLAB (The Mathworks, Natick, MA, USA) that allows the extraction of parameters describing the burst shape. The parameters extracted using Multiwell analyser include the mean firing rate (MFR), which is the average total number of spikes detected per electrode over time (spikes/second in Hz). The MFR is averaged per well for all active electrodes. The PRS is calculated from spikes that were not included in the burst, nor network burst. We detected bursts based on the maximum inter-spike interval (ISI) to start or end a burst (i.e., for the whole spike train, with a maximum ISI of 30 ms). If the ISI is shorter than 30 ms spikes were included in the burst, if the ISI is larger than 30 ms the burst ends. All bursts that were less than 65 ms apart were merged. All bursts that have a duration of less than 50 ms or have less than 4 spikes were removed from the detection^84^. Network bursts were detected when a burst occurs in more than 80% of the active channels. The NBR is calculated as network burst/minute and the NBD as duration in ms. Not active wells (i.e. MFR < 0.16 Hz in at least 3 channels to be called active), wells that had no NB at DIV 28, wells where network bursts were detected in less than 50% of the channels and wells where the firing rate decreased over development were rigorously discarded. Discriminant function analysis based on parameters describing neuronal network activity were performed in SPSS (IBM Corporation, Armonk, NY, USA).

### Single cell electrophysiology

Coverslips were placed in the recording chamber of the electrophysiological setup, continuously perfused with oxygenated (95% O_2_/ 5% CO_2_) ACSF at 32°C containing (in mM) 124 NaCl, 1.25 NaH_2_PO_4_, 3 KCl, 26 NaHCO_3_, 11 Glucose, 2 CaCl_2_, 1 MgCl_2_. Patch pipettes with filament (6-8 MΩ) were pulled from borosilicate glass (Science Products GmbH, Hofheim, Germany) using a Narishige PC-10 micropipette puller (Narishige, London, UK). For all recordings of intrinsic properties and spontaneous activity, a potassium-based solution containing (in mM) 130 K-Gluconate, 5 KCl, 10 HEPES, 2.5 MgCl_2_, 4 Na_2_-ATP, 0.4 Na_3_-ATP, 10 Na-phosphocreatine, 0.6 EGTA (pH 7.2 and 290 mOsmol) was used. vRMP was measured immediately after generation of a whole-cell configuration. Further analysis of active and passive membrane properties was conducted at a holding potential of −60 mV. Passive membrane properties were determined via voltage step of −10 mV. Active intrinsic properties were measured with a stepwise current injection protocol. Spontaneous activity was measured at either −60 mV (sPSCs, drug free) or +10 mV (GABAergic sPSCs, 100 μM CNQX, Tocris). Activity was recorded at DIV 28, 35 and 49. Cells were visualized with an Olympus BX51WI upright microscope (Olympus Life Science, PA, USA), equipped with a DAGE-MTI IR-1000E (DAGE-MTI, IN, USA) camera) and a CoolLED PE-200 LED system (Scientifica, Sussex, UK) for fluorescent identification. A Digidata 1440A digitizer and a Multiclamp 700B amplifier (Molecular Devices) were used for data acquisition. Sampling rate was set at 20 kHz and a lowpass 1 kHz filter was used during recording. Recordings were not corrected for liquid junction potential (± 10 mV). Recordings were discarded if series resistance reached >25 MΩ or dropped below a 10:0 ratio of membrane resistance to series resistance. Intrinsic electrophysiological properties were analysed using Clampfit 10.7 (molecular devices, CA, USA), and sPSCs were analysed using MiniAnalysis 6.0.2 (Synaptosoft Inc, GA, USA) as previously described^77^.

For the determination of decay times, GABAergic events were isolated in neurons at DIV 49 by bath application of CNQX. This decay time was then compared to the decay time of glutamatergic events recorded in the presence of PTX. We determined that a cut-off of 3.8 ms (supplementary figure 2s) could to a high degree of confidence separate glutamatergic and GABAergic events in other data. This cut-off was then used to separate glutamatergic and GABAergic events during development.

### Pharmacological Experiments

Picrotoxin (PTX) was prepared fresh into concentrated stocks and stored frozen at −20°C (PTX, 50 mM in ETOH (MEA) or 100 mM in DMSO (single-cell recordings), Tocris Cat No 1128). For all experiments on MEAs an aliquot of the concentrated stock was first diluted 1:2.5 in room temperature DPBS and vortexed briefly. Then, 2.5 μl working dilution was added directly to wells on the MEA and mixing was primarily through diffusion into the (500 μl) cell culture medium to reach a 100 μM concentration. ETOH was used as vehicle and similarly diluted as the PTX. For all single cell experiments, PTX was directly diluted 1000 x in artificial cerebrospinal fluid (ACSF). The DMSO concentration in the ACSF was always ≤0.05% v/v. All experiments were performed at 37°C.

### Cell adhesion assay

The cell aggregation assay was performed as described previously^60^. HEK293T cells were transfected with indicated constructs via calcium phosphate transfection when they reached a confluency of 50%. To assess the rate of transfection, additional vectors encoding GFP or mCherry were used to mark the transfected cells. At least 26h after transfection the rate of successfully transfected cells was determined. In case the transfection rate was above 75% aggregation assays were performed.

Cells were trypsinized and collected by centrifugation for 5 min at 4°C and 1000 rpm,and washed once with serum-free medium, before being resuspended by pipetting in Hank’s Balanced Saline Solution (HBSS) (55 mM NaCl, 40 mM KCl, 15 mM MgSO_4_, 10mM CaCl_2_, 20mM glucose, 50 mM sucrose, 2 mg/ml bovine serum albumin, and 20 mM Tricine, pH 6.95). Cell concentration was determined by counting with a hemocytometer. HBSS was used to dilute the cell suspension to a final concentration of 1.2x 10^6^ cells/ml for single line experiments, or 6x 10^5^ hhen two different cell lines were incubated. 1 ml cell suspension was filled into 1.5 ml Eppendorf tubes and incubated at 4 °C under gentle agitation for 1 hour. Aggregation was quantified by counting the cells with a hemocytometer. Cell aggregation was plotted as the ratio *T*0/*T60*, where *T*0 is the total number of cellular particles before incubation, and *T60* is the total number of cellular particles after 1 hour incubation, with cellular aggregates counting as single particles.

### Statistics

The statistical analysis for all experiments was performed using GraphPad Prism 8 (GraphPad Software, Inc., CA, USA). We ensured normal distribution using a Kolmogorov-Smirnov normality test. To determine statistical significance for the different experimental conditions *p*-values <0.05 were considered to be significant. Statistical analysis of electrophysiological data in Figure 1 and 4d were performed with one-way ANOVA and post hoc Tukey (normal distribution) or Kruskal-Wallis ANOVA with post hoc Dunn’s (not normally distributed) correction for multiple testing. Statistical analysis over development (Figure 3) were performed with two-ways ANOVA and Post-hoc Bonferroni (normal distribution) or a Mixed effects analysis and post hoc Dunn’s (not normally distributed) correction for multiple testing (depending on normal distribution). When comparing means of two variables at one individual timepoint we analysed significance between groups by means of a two sided paired T-test (when using paired data; Figure 4h-k, Supplementary figure 3b, k-m) or Mann-Whitney-U-Test (unpaired data; Figure 2c, f, 4g, 5i, l, m and Supplementary figure 3j, 4c, e), and if applicable, corrected post hoc for multiple testing using the Bonferroni method. Statistics on histograms was performed using Multiple t-test on bins using Holm-Sidak method (figure 4h-k, 5j). Data are presented as Mean ± standard error of the mean (SEM). Means and *p*-values are reported in Supplementary Tables 1–9.

### Cell line information and hiPSC generation

In this study we used in total 3 control hIPSC lines, Control #1, Control #2 and Control #3. All hiPS cells used in this study were obtained from reprogrammed fibroblasts. Control line 1 was obtained from a healthy 30-year-old female, and reprogrammed via episomal reprogramming^85^. Control line 2 was derived from a healthy 51-year-old male and reprogrammed via a non-integrating Sendai virus by KULSTEM (Leuven, Belgium). Control line 3 was obtained from a healthy male, and reprogrammed via retroviral vectors expressing four transcription factors: Oct4, Sox2, Klf4, and cMyc. Generated clones (at least two per patient line) were selected and tested for pluripotency and genomic integrity based on single nucleotide polymorphism (SNP) arrays^75^. HiPSCs were cultured on Matrigel (Corning, #356237) in E8 flex (Thermo Fisher Scientific) supplemented with primocin (0.1 μg/ml, Invivogen) and low puromycin (0.5 μg/ml, to select for rtTA positive cells) and G418 concentrations (50 μg/ml, to select for *Ngn2* or *Ascl1* positive cells) at 37°C/5% CO_2_. Medium was refreshed every 2-3 days and cells were passaged twice a week using an enzyme-free dissociation reagent (ReLeSR, Stem Cell Technologies).

### Animals

The rodent astrocytes presented in this study were harvested embryonic (E18) rat brains (Wistar Wu) as previously described^75, 77, 86^. All experiments on animals were carried out in accordance with the approved animal care and use guidelines of the Animal Care Committee, Radboud University Medical Centre, the Netherlands, (RU-DEC-2011-021, protocol number: 77073).

### Compound application

The GABA reversal was measured using a cesium-based intracellular solution containing (in mM) 115 CsMeSO_3_, 20 CsCl, 10 HEPES, 2.5 MgCl_2_, 4 Na_2_ATP, 0.4 Na_3_GTP, 10 sodium phosphocreatine, 0.6 EGTA (pH 7.2, mOsmol 290). For sucrose application, cells were recorded with the KCl based solution described before.

GABA (10 mM dissolved in ACSF) was applied locally at a distance of 10-20 μm from the soma of the patched excitatory neuron using a PDES-2DX pressure ejection system (NPI, Tamm, Germany). Micropipettes used for compound application had a resistance of 3-5 MΩ. Injection pressure was set at 7psi/0.5 bar and injection duration was set to 100 ms. Analysis of peak response and reversal potential was conducted using Clampfit 10.7.

### Immunocytochemistry

Cells were fixed and stained as described before^75^. All antibodies are listed in Supplementary table 11. Neurons were generally fixated at DIV 49, and at DIV 35 and DIV 49 for membrane expression of NKCC1 and KCC2. When membrane expression of NKCC1 and KCC2 was examined, coverslips were not permeabilised. We imaged at a 20x magnification to count the number of GABAergic subtypes and at a 63x magnification for all other measures using the Zeiss Axio Imager Z1 equipped with apotome. All conditions within a batch were acquired with the same settings in order to compare signal intensities between different experimental conditions. Fluorescent signals were quantified using FIJI software. The intensity of NKCC1 or KCC2 expression on the cell membrane was calculated by: integrated density – (Area of selected cell X Mean fluorescence of background readings). The number of synaptic puncta was determined per individual cell via manual counting and divided by the dendritic length of the dendrite. VGAT puncta intensity was determined using particle analysis in the FIJI software.

### Quantification of mRNA by RT-qPCR

RNA samples were isolated using Nucleospin RNA isolation kit (Machery Nagel, 740955.250) according to the manufacturer’s instructions. RNA samples were converted into cDNA by iScript cDNA synthesis kit (BIO-RAD, 1708891). CDNA products were cleaned up using the Nucleospin Gel and PCR clean-up kit (Machery Nagel, 740609.250). Human-specific primers were designed with Primer3plus (http://www.bioinformatics.nl/cgi-bin/primer3plus/primer3plus.cgi) and IDT PrimerQuest (https://eu.idtdna.com) tools, respectively. Primer sequences are given in supplementary table 12. QPCRs were performed in the Quantstudio 3 apparatus (Thermo Fisher Scientific) with GoTaq qPCR master mix 2X with SYBR Green (Promega, A600A) according to the manufacturer’s protocol. The qPCR program was designed as following: After an initial denaturation step at 95°C for 10 min, PCR amplifications proceeded for 40 cycles of 95°C for 15 s and 60°C for 30 s and followed by a melting curve. All samples were analysed in duplicate in the same run, placed in adjacent wells. The arithmetic mean of the Ct values of the technical replicas was used for calculations. Relative mRNA expression levels were calculated using the 2^-ΔΔCt method with standardization to housekeeping genes^87^.

### RNA-sequencing

RNA was isolated from three biological replicates of E/I networks composed of *Ngn2* #2 and *Ascl1* #1 (DIV 49) with the *Quick*-RNA Microprep kit (Zymo Research, R1051) according to manufacturer’s instructions. RNA quality was checked using Agilent’s Tapestation system (RNA High Sensitivity ScreenTape and Reagents, 5067-5579/80). RIN values ranged between 7.5 – 8.3. RNA-sequencing (RNA-seq) library preparation was performed using a published single-cell RNA-seq protocol from Cao et al. 2017^88^ which was adapted for bulk RNA-seq experiments. For each sample, 10 ng total RNA (in 0.65 μL) was mixed with 0.1 μL dNTP mix (10 mM each) (Invitrogen, 10297018), 0.15 μL ERCC RNA Spike-In Mix (100.000x diluted) (Thermo Scientific, 4456740), 0.15 μL nuclease-free water (NF H_2_O) and 0.4 μL anchored oligo-dT (2.5 μM) primer(5ʹ-ACGACGCTCTTCCGATCTNNNNNNNN[10bp index]TTTTTTTTTTTTTTTTTTTTTTTTTTTTTTVN-3ʹ, where “N” is any base and “V” is either “A”, “C” or “G”; IDT) in a tube containing 7 μL Vapor-Lock (Qiagen, 981611) to prevent evaporation. Each sample was incubated for 5 min at 65°C and directly placed on ice. First strand reaction mix was added, consisting of 0.4 μL Maxima RT buffer (5X) (Thermo Scientific, EP0751), 0.05 μL RNasin Plus (Promega, N2611) and 0.1 μL Maxima H Minus Reverse Transcriptase (Thermo Scientific, EP0751). Reverse transcription was performed by incubating the samples at 50°C for 30 min and terminated by heating at 85°C for 5 min. For second strand synthesis, 2 μL RT product was mixed with 7.7 μL NF H_2_O, 2.5 μL Second Strand Buffer (Invitrogen, 10812014), 0.25 μL dNTP mix (10 mM each), 0.35 μL DNA polymerase I (*E. coli*) (NEB, M0209), 0.09 μL DNA ligase (*E. coli*) (NEB, M0205) and 0.09 μL Ambion RNase H (*E. coli*) (Invitrogen, AM2293). Second strand synthesis was performed by incubating samples at 16°C for 150 min, followed by 75°C for 20 min. Next, 0.5 μL Exonuclease I (NEB, M0293) was added per sample and incubated at 37°C for 60 min. cDNA samples were pooled per sets of 6-8 samples, Vapor-Lock was removed and samples were added up with NF H_2_O to a total volume of 107.6 μL. Each pool of samples was then purified using 79 μL beads buffer (20% PEG-8000 in 2.5 M NaCL, final concentrations) and 50 μL Ampure XP Beads (Beckman Coulter, A63881), and eluted in 7 μL NF H_2_O.

Tagmentation was performed per pool by adding 3 μL double-stranded cDNA sample to 5.5 μL Nextera TD buffer (Illumina, 15027866), 2.5 μL NF H_2_O and 1.0 μL TDE1 Enzyme (Illumina, 15027865), which was incubated at 55°C for 5 min. Samples were directly placed on ice for at least 3 min. The reaction was terminated by adding 12 μL Buffer PB (QiaQuick, 19066) and incubating for 5 min at room temperature. Samples were purified using 48 μL Ampure XP beads (Beckman Coulter, A63881) and eluted in 10 μL NF H_2_O. Next, each sample was mixed with 2 μL P5 primer (10 μM), (5ʹ-AATGATACGGCGACCACCGAGATCTACAC [i5]ACACTCTTTCCCTACACGACGC TCTTCCGATCT-3ʹ;IDT), 2 μL P7 primer (10 μM) (5ʹ-CAAGCAGAAGACGGCATACGAG AT[i7] GTCTCGTGGGCTCGG-3ʹ; IDT) and 20 μL NEBNext High-Fidelity 2X PCR Master Mix (NEB, M0541). Amplification was performed using the following program: 72°C for 5 min, 98°C for 30 sec, 15 cycles of (98°C for 10 sec, 66°C for 30 sec, 72°C for 1 min) and a final step at 72°C for 5 min. Samples were purified using 32 μL Ampure XP beads (Beckman Coulter, A63881) and eluted in 12 μL NF H_2_O. Libraries were visualized by electrophoresis on a 1% agarose and 1X TAE gel containing 0.3 μg/mL ethidium bromide (Invitrogen, 15585011). Gel extraction was performed to select for products between 200 – 1000 bp using the Nucleospin Gel and PCR Clean-up kit (Macherey-Nagel, 740609). Samples were eluted in 11 μL NF H_2_O. cDNA concentrations were measured by Qubit dsDNA HS Assay kit (Invitrogen, Q32854). Product size distributions were visualized using Agilent’s Tapestation system (D5000 ScreenTape and Reagents, 5067-5588/9). Libraries were sequenced on the NextSeq 500 platform (Illumina) using a V2 75 cycle kit (Read 1: 18 cycles, Read 2: 52 cycles, Index 1: 10 cycles).

### Pre-processing of RNA-seq data

Base calls were converted to fastq format and demultiplexed using Illumina’s bcl2fastq conversion software (v2.16.0.10) tolerating one mismatch per library barcode. Reads were filtered for valid unique molecular identifier (UMI) and sample barcode, tolerating one mismatch per barcode. Trimming of adapter sequences and over-represented sequences was performed using Trimmomatic (version 0.33)^89^. Trimmed reads were mapped to a combined human (GRCh38.p12) and rat (Rnor_6.0) reference genome to separate reads belonging to the human iNeurons from reads originating from the rat astrocytes. Mapping was performed using STAR^90^ (version 2.5.1b) with default settings (--runThreadN 1, --outReadsUnmapped None, --outFilterType Normal, --outFilterScoreMin 0, --outFilterMultimapNmax 10, -- outFilterMismatchNmax 10, --alignIntronMin 21, --alignIntronMax 0, --alignMatesGapMax 0,--alignSJoverhangMin 5, --alignSJDBoverhangMin 3, --sjdbOverhang 100). Uniquely mapped reads (mapping quality of 255) were extracted and read duplicates were removed using the UMI-tools software package^91^. Raw reads from BAM files were further processed to generate count matrices with HTSeq^92^ (version 0.9.1) using reference transcriptome Gencode GRCh38.p12 (release 29, Ensembl 94). Raw counts were transformed to log-transformed counts per million (logCPM) using edgeR (R package). A count table with raw counts/(log)cpm values can be found in supplementary table 13, and are deposited in GEO with the accession code GSE144197.

### Western blot

Precleared HEK293T cells lysates co-transfected with Myc-tagged *Cdh13*, GFP-tagged *Itgβ3* and myc-tagged *GABAAα1* for 24 hours were incubated with mouse IgG (eBioscience), anti-myc (Abcam) or anti-GFP (Invitrogen) antibodies (2.5 μg/ml lysate) at 4°C for 1h. Dynabeads® Protein G (50 μl per sample, Invitrogen) were then added and incubated for 1h at 4°C under gentle rotation. Samples were washed three times with lysis buffer (20 mM Tris, 150 mM NaCl, 2 mM EDTA, 1 mM DTT and protease inhibitor, pH7.4). Co-immunoprecipitated proteins were detected by Western blotting. Samples were fractionated on 4%-15% precast SDS-PAGE gels (Bio-Rad) and then blotted on a nitrocellulose membrane (Bio-Rad). Membranes were subsequently incubated for 1h at room temperature in 5% milk in PBS-Tween to prevent non-specific binding. Primary antibodies used were: anti-myc (Abcam), anti-GFP (Invitrogen) or anti-Y-TUBULIN (Sigma) as loading control. Secondary antibodies used were: goat anti-mouse HRP (Jackson Immuno Research Laboratories) and goat anti-rabbit (Invitrogen). Images were acquired with ChemiDocTM imaging system (Bio-Rad).

## Acknowledgements

We thank dr. J. Ladewig and prof. Dr. O. Brüstle for providing the *Ngn2* and *Ascl1* lentiviral construct.

## Author contributions

B.M, D.S and N.N.K conceived and designed all the experiments. N.N.K, D.S and H.v.B supervised the study. B.M, J.v.R, S.W, E.v.H, K.L, J.B, A.V, S.J, J.K, T.K.G, C.S. A.O performed all experiments. H.v.B, and M.S provided resources. B.M, J.v.R, J.B, S.W, A.V, A.A, M.F, D.S, N.N.K performed data analysis. B.M, J.v.R, D.S and N.N.K wrote the manuscript. H.v.B, M.F, M.S, A.V, E.v.H and J.K edited the manuscript.

## Competing interests

The authors declare no competing interests.

Correspondence and requests for materials should be addressed to

## Supplementary Figures

**Supplementary figure 1.**
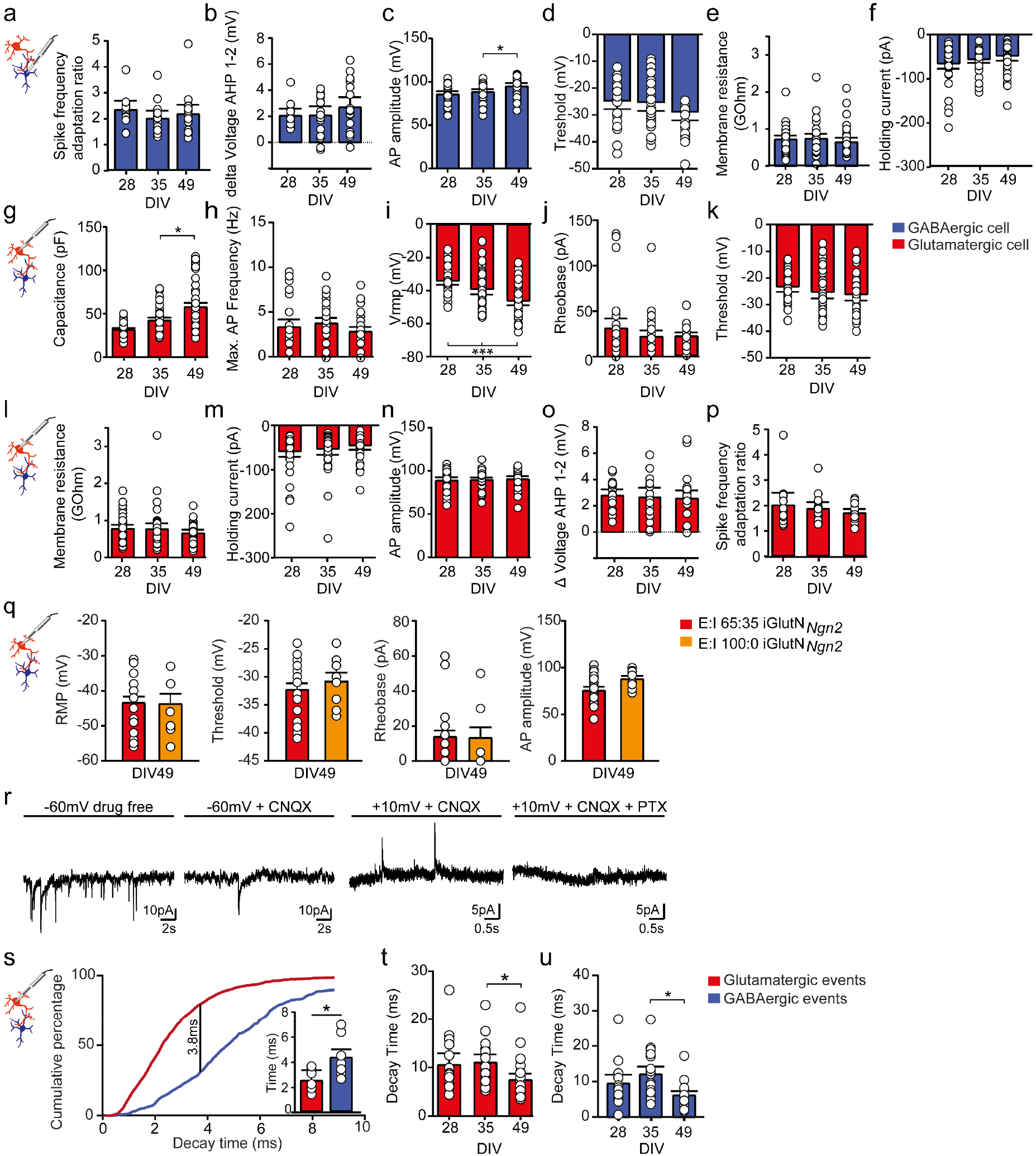
iGLU_Ngn2_ and iGABA_A-FSK_ mature over development. (**a-f**) Passive and active intrinsic properties recorded from iGABA_A-FSK_ in an E:I 65:35 culture (Sample size for DIV 28 n=39, DIV 35 n=38, DIV 49 n=41 cells from 3 batches). (**g-p**) Passive and active intrinsic properties recorded from iGLU_Ngn2_ in an E:I culture (Sample size for DIV 28 n=42, DIV 35 n=40, DIV 49 n=44 recorded cells from 3 individual neuronal preparations). (**q**) Intrinsic properties recorded from iGLU_Ngn2_ in E:I 65:35 (red) versus iGLU_Ngn2_ in an E:I 100:0 (yellow) only culture (iGLU_Ngn2_ in E:I 65:35 networks n=23 and iGLU_Ngn2_ in E:I 100:0 networks n=8 recorded cells from 2 individual neuronal preparations). (**r**) Representative traces of spontaneous network activity (i.e. Glutamatergic and GABAergic sPSCs) under drug free conditions (panel 1), when AMPA receptors are blocked with 6-cyano-7-nitroquinoxaline-2,3-dione (CNQX, GABAergic sPSCs in panel 2 and 3), or when AMPA and GABA receptors are blocked with CNQX and Picrotoxin (PTX, panel 4). (**s**) Cumulative plot of decay time of either GABAergic or glutamatergic events. Largest difference at 3.8 ms explains 78% percent of variance. (**s-u**) Average decay time calculated from spontaneous activity that was split at decay time of 3.8 ms to distinguish the (**t**) Glutamatergic and (**u**) GABAergic events. DIV: Days in vitro. All data represent means ± SEM. * p < 0.05, *** p < 0.001. Mann-Whitney test with post hoc Bonferroni correction for multiple testing was performed between DIVs.

**Supplementary figure 2.**
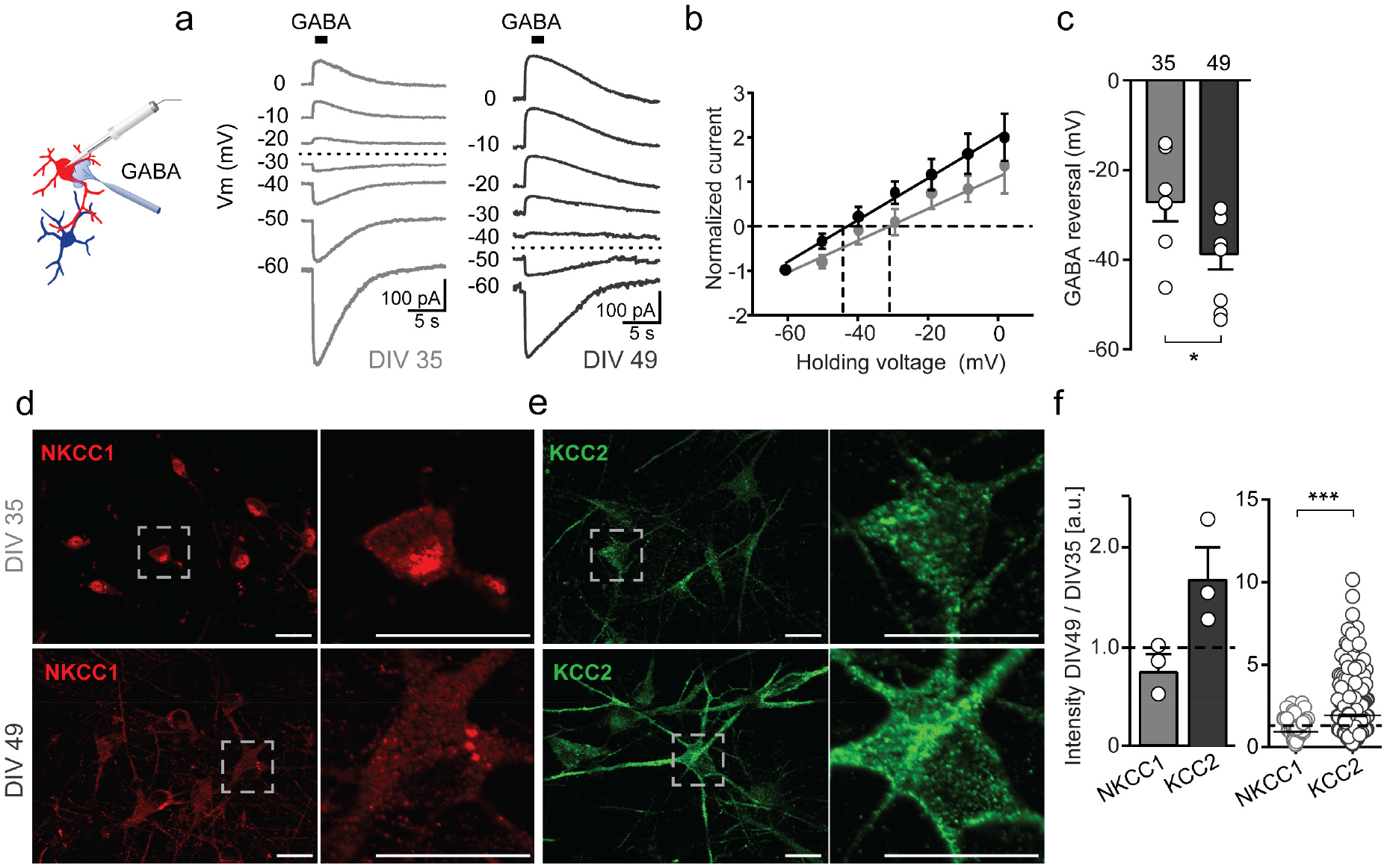
iGABA_A+FSK_ become functionally inhibitory at DIV49. (**a**) Representative traces showing GABA-evoked currents of iGLU_Ngn2_ under various holding potentials at DIV 35 and −49. (**b**) Quantification of GABA-evoked responses. Dashed lines represent GABA reversal potential. (**c**) Quantified results of the reversal potential at DIV 35 and DIV 49 (DIV 35 n=7 and DIV 49 n=10 cells from 2 neuronal preparations). (**d-e**) Representative (**d**) NKCC1 and (**e**) KCC2 immunostaining in E/I networks at DIV 35 and −49. (**f**) NKCC1 and KCC2 intensity measurements at DIV 49 normalised to the expression levels of DIV 35 (dashed line). Left panel: each data point represents one neuronal preparation. Right panel: Each data point represents the normalised intensity of one cell (NKCC1 DIV 35 n=206; NKCC1 DIV 49 n=153; KCC2 DIV 35 n=256; KCC2 DIV 49 n=237 cells analysed from 3 different neuronal preparations). DIV: Days in vitro. All data represent means ± SEM. * p < 0.05; *** p < 0.001 (Mann-Whitney test with post hoc Bonferroni correction for multiple testing was performed between DIVs). Scale bar 10 μM.

**Supplementary Figure 3.**
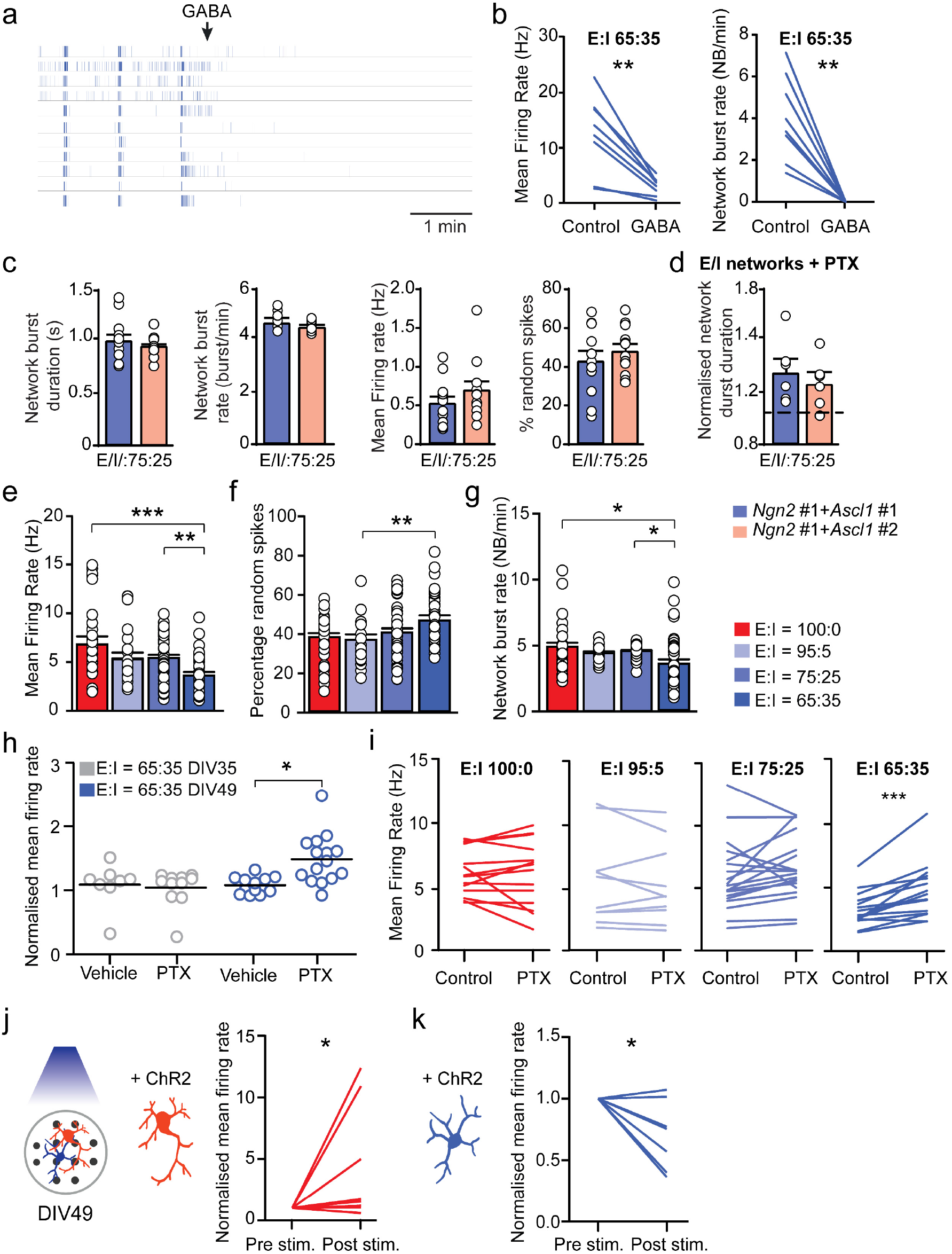
Functional GABAergic modulation in E/I networks in depending on the hyperpolarizing GABA shift and scalable to the percentage iGABA_A-FSK_ present in the network. (**a**) Representative raster plot of a 7 minute recording of an E:I 65:35 network treated with 100 μM Gamma-Aminobutyric acid (GABA). (**b**) Paired analysis of the average mean firing rate (MFR, left) and network burst rate (NBR, right) before and after treatment with GABA in E:I 65:35 networks at DIV 49 (Sample size n for E:I 65:35 networks n=9 individual wells, paired T-test was performed between pre and post conditions). (**c-d**) Comparison between E/I networks composed of two different *Ascl1* stable lines (i.e. E:I 75:25 networks from *Ngn2* #1 and *Ascl1* #1 and *Ngn2* #1 and *Ascl1* #2) at DIV 49 at the level of (**c**) spontaneous network activity and (**d**) response of the network burst duration to picrotoxin (PTX) treatment (Sample size n for *Ngn2* #1 and *Ascl1* #1 n=12 and *Ngn2* #1 and *Ascl1* #2 n=15 individual cultures from 2 individual neuronal preparations). (**e-g**) Quantifications of (**e**) MFR, (**f**) percentage of random spikes and (**g**) NBR from E:I 100:0, 95:5, 75:25 and 65:35 networks (Sample size for E:I 100:0 n=29, E:I 95:5 n=20, E:I 75:25 n=38 and E:I 65:35 n=38 individual wells, Kruskal-Wallis Two way ANOVA was performed and corrected for multiple testing using Dunn’s method). (**h**) MFR of vehicle or 100 μM PTX treated E:I 65:35 networks, normalised to their respective baseline recording (Sample size for DIV 35 + vehicle n= 8; DIV 35 + PTX n=11; DIV 49 + vehicle n=12 and DIV 49 + PTX n=15 individual wells, Mann-Whitney T-test was performed, *p* values were corrected for multiple testing using Bonferroni method). (**i**) Paired analysis of the average MFR before and after treatment with 100 μM PTX (Sample size for 100:0 cultures n=15, 95:5 n=10, 75:25 n=19 and 65:35 n=15 individual wells, paired T-test was performed between pre and post conditions). (**j-k**) MFR of 65:35 networks upon cell-type specific optogenetic activation of iGLU_Ngn2_ (**j**) or iGABA_A-FSK_ (**k**) neurons respectively at DIV 49. Pre stimulation condition represents the MFR normalized to 50 ms pre-stimulation baseline activity. Post stimulation condition represents the activity in a window between 10-30 ms after stimulus onset. MFR was normalised to pre-stimulation condition (sample size n=7 individual wells for both conditions, paired T-test (Wilcoxon rank-sum) was performed between pre and post conditions). DIV: Days in vitro. All data represent means ± SEM. * p < 0.05; ** p < 0.01; *** p < 0.001.

**Supplementary figure 4.**
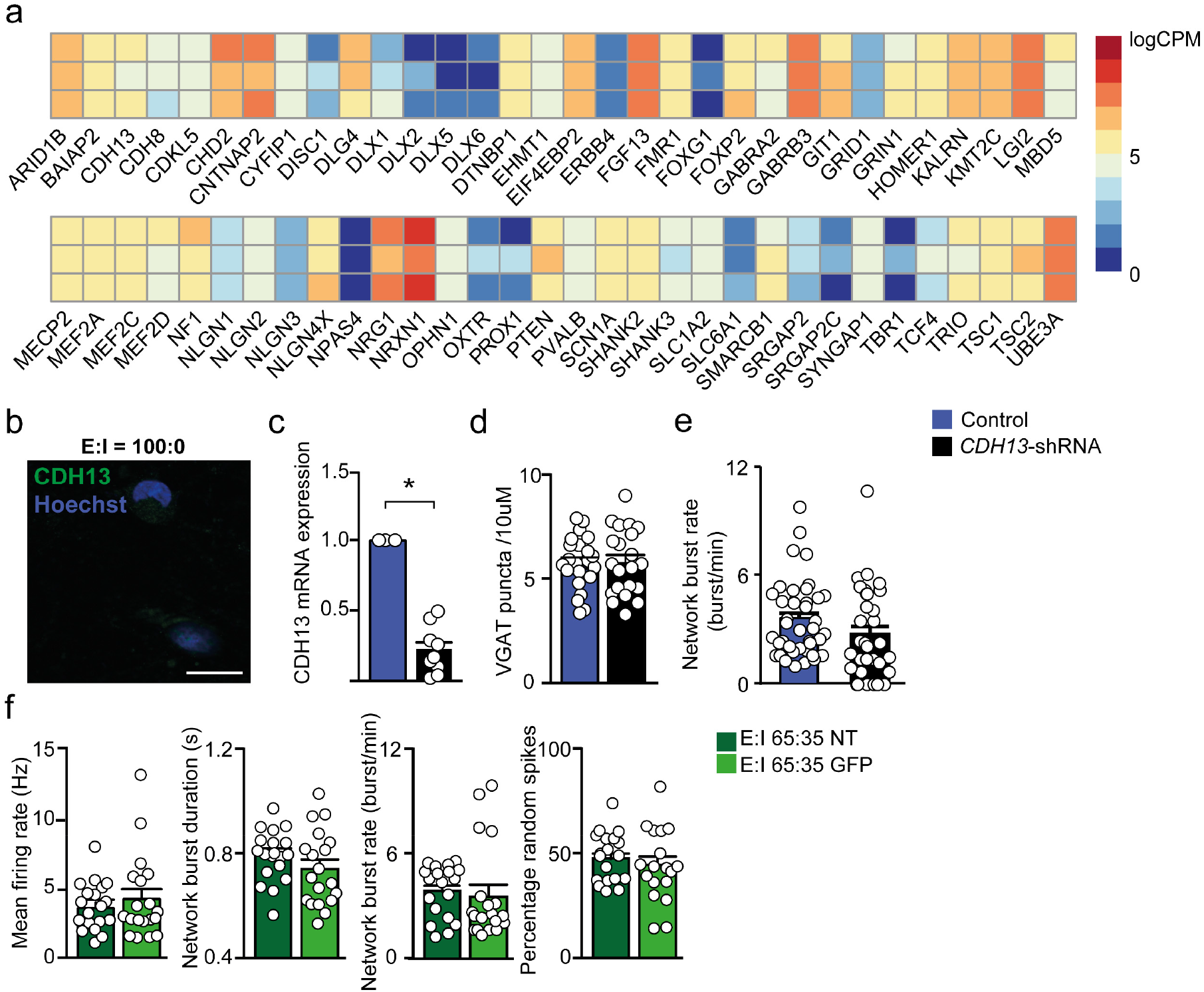
E/I cultures pose a valid model to study cell-type specific interactions of NDD genes to network dysfunction; an example for CDH13-deficiency. (**a**) Bulk RNA sequencing analysis of E:I 65:35 networks at DIV 49 (3 biological replicates). (**b**) CDH13 expression in iGLU_Ngn2_ neurons at DIV 49. (**c**) CDH13 mRNA expression following shRNA mediated knockdown. (**d**) Total number of VGAT positive presynapses in control and CDH13-deficient networks at DIV 49 (control n=23, CDH13-deficient n=21, analysed cells from 3 individual neuronal preparations). (**e**) Network burst rate of control and CDH13-deficient networks at DIV 49 (Sample size n for 65:35 control n= 38 and 65:35 CDH13-deficient wells n= 33 individual wells from 3 neuronal preparations. Mann-Whitney test with post hoc Bonferroni correction for multiple testing was performed, *p*=0.065). (**f**) Network activity compared between non-treated 65:35 controls (NT) and GFP scrambled shRNA controls (GFP) at DIV 49 (Sample size n for E:I 65:35 control n= 20 and E:I 65:35 GFP scrambled shRNA n= 20 individual wells from 3 neuronal preparations). DIV: Days in vitro. All data represent means ± SEM. * p < 0.05.

## Supplementary tables

**Supplementary table 1.** Statistics of intrinsic properties from **iGABA_A-FSK_** in **Figure 1h, i, n** and **Supplementary figure 2c, s**. Sample size for intrinsic properties at DIV 28 n=39, DIV 35 n=38, DIV 49 n=41 recorded cells from 3 neuronal preparations. Sample size of correlated synaptic inputs at DIV 28 n=55, DIV 35 n=38, DIV 49 n=42 recorded cells from 3 neuronal preparations. All data represent means ± SEM. Two-way ANOVA with Tukey correction for multiple testing was used to compare between DIVs. DIV= Days in vitro.

| Panel |  | DIV | Mean | SEM | p-value | Comparison to |
| --- | --- | --- | --- | --- | --- | --- |
| **Fig. 1h** | Rmp | 28 | -32.3 | 1.5 |  |  |
|  |  | 35 | -38.3 | 1.6 | <0.001 | DIV 49 |
|  |  | 49 | -50.2 | 1.3 | <0.001 | DIV 28, DIV 35 |
| **Fig. 1i** | Capacitance | 28 | 37.6 | 1.7 |  |  |
|  |  | 35 | 43.1 | 2.2 | <0.05 | DIV 49 |
|  |  | 49 | 57.8 | 3.5 | <0.001 | DIV 28 |
| **Fig. 1n** | Correlated synaptic input | 28 | 0.19 | 0.08 |  |  |
|  |  | 35 | 0.24 | 0.08 | <0.001 | DIV 49 |
|  |  | 49 | 0.65 | 0.11 | <0.001 | DIV 28, DIV 35 |
| **Sup. Fig. 2c** | Ap amplitude | 28 | 85.5 | 2.0 |  |  |
|  |  | 35 | 88.1 | 1.8 |  |  |
|  |  | 49 | 94.6 | 2.1 | <0.05 | DIV 35 |
| **Sup. Fig. 2s** | Decay time | 49 | 2.39 | 0.32 |  |  |
|  |  | 49 | 4.23 | 0.65 | <0.05 |  |
| **Sup. Fig. 2t** | Decay time | 28 | 8.81 | 2.0 |  |  |
|  |  | 35 | 12.1 | 1.68 |  |  |
|  |  | 49 | 6.15 | 0.78 | <0.05 | DIV35 |
| **Sup. Fig. 2u** | Decay time | 28 | 9.45 | 2.35 |  |  |
|  |  | 35 | 12.05 | 2.01 |  |  |
|  |  | 49 | 6.12 | 0.90 | <0.05 | DIV35 |

**Supplementary table 2.**
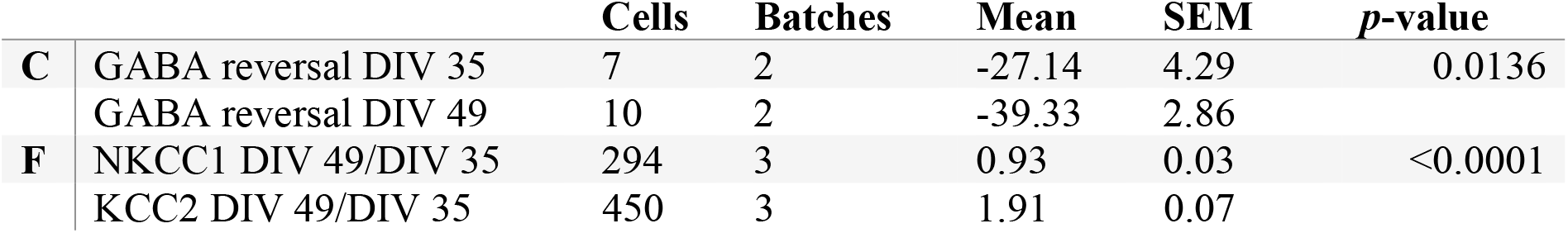
Statistics of **Figure 2**. All data represent means ± SEM. * p < 0.05; ** p < 0.01; *** p < 0.001. Batch number indicates the amount of neuronal preparations. Mann-Whitney test with post hoc Bonferroni correction for multiple testing was performed between DIVs. DIV= Days in vitro.

**Supplementary table 3.** Statistics of **supplementary figure 2g-j**. All data represent means ± SEM. * p < 0.05; ** p < 0.01; *** p < 0.001. Mixed model Two way ANOVA was performed between DIVs, p values were corrected for multiple comparisons using Sidak’s. NBD= Network burst duration, NBR= Network burst rate, MFR= Mean firing rate, PRS= percentage of random spikes, n= number of wells, DIV= Days in vitro.

|  | DIV | E:I=100:0 |  |  | E:I=65:35 |  |  | *p*-value E:I<br>65:35 vs 100:0 | *p*-value<br>development |
| --- | --- | --- | --- | --- | --- | --- | --- | --- | --- |
|  |  | Mean | SEM | n | Mean | SEM | n |  |  |
| **NBD (MS)** | 35 | 1228.60 | 81.38 | 25 | 1059.57 | 41.74 | 36 | 0.2026 |  |
|  | 42 | 1108.37 | 57.01 | 30 | 853.13 | 24.38 | 39 | 0.0006 |  |
|  | 49 | 1114.40 | 31.42 | 29 | 750.19 | 18.46 | 37 | <0.0001 | <0.0001 |
| **NBR (B/MIN)** | 35 | 3.27 | 0.24 | 25 | 2.40 | 0.16 | 40 | 0.0143 |  |
|  | 42 | 4.45 | 0.27 | 30 | 2.62 | 0.21 | 39 | <0.0001 |  |
|  | 49 | 4.90 | 0.33 | 29 | 3.562 | 0.32 | 37 | 0.0150 | <0.0001 |
| **MFR (HZ)** | 35 | 3.62 | 0.54 | 25 | 2.322 | 0.27 | 40 | 0.1053 |  |
|  | 42 | 6.43 | 0.85 | 30 | 2.98 | 0.28 | 39 | 0.0014 |  |
|  | 49 | 7.06 | 0.67 | 29 | 3.68 | 0.31 | 38 | <0.0001 | <0.0001 |
| **PRS (%)** | 35 | 53.37 | 3.86 | 25 | 55.87 | 2.83 | 40 | 0.9382 |  |
|  | 42 | 44.26 | 3.06 | 30 | 53.33 | 2.38 | 39 | 0.0664 |  |
|  | 49 | 37.54 | 2.43 | 29 | 47.52 | 2.18 | 38 | 0.0100 | <0.0001 |

**Supplementary table 4.** Statistics of **Figure 3d** and **supplementary figure 3e-g**. All data represent means ± SEM. * p < 0.05; ** p < 0.01; *** p < 0.001. Kruskal Wallis one way ANOVA was performed between ratio’s, p values were corrected for multiple comparisons using Dunn’s. Multiple comparison statistics are mentioned last column, compared ratio’s were split by ‘/’. Other comparisons were non-significant. M=Mean, n= number of wells, NBD= Network burst duration, NBR= Network burst rate, MFR= Mean firing rate, PRS= percentage of random spikes.

|  | E:I=100:0 |  |  | E:I=95:5 |  |  | E:I=75:25 |  |  | E:I=65:35 |  |  | *p*-value mult. comp. |
| --- | --- | --- | --- | --- | --- | --- | --- | --- | --- | --- | --- | --- | --- |
|  | M | SEM | n | M | SEM | n | M | SEM | n | M | SEM | n |  |
| **NBD (MS)** | 1114 | 31.4 | 30 | 1045 | 56.0 | 19 | 901.8 | 26.9 | 42 | 751.2 | 17.5 | 39 | 100:0/75:25=0.0006 |
|  |  |  |  |  |  |  |  |  |  |  |  |  | 100:0/65:35=<0.0001 |
|  |  |  |  |  |  |  |  |  |  |  |  |  | 95:5/65:35=0.0002 |
|  |  |  |  |  |  |  |  |  |  |  |  |  | 75:25/65:35=0.0015 |
| **NBR (b/min)** | 4.9 | 0.3 | 30 | 4.5 | 0.1 | 19 | 4.6 | 0.1 | 42 | 3.6 | 0.3 | 39 | 100:0/65:35=0.0220 |
|  |  |  |  |  |  |  |  |  |  |  |  |  | 75:25/65:35=0.0209 |
| **MFR (Hz)** | 7.1 | 0.7 | 30 | 5.4 | 0.6 | 19 | 5.3 | 0.4 | 42 | 3.7 | 0.3 | 39 | 100:0/65:35=<0.001 |
|  |  |  |  |  |  |  |  |  |  |  |  |  | 75:25/65:35=0.0097 |
| **PRS (%)** | 37.5 | 2.4 | 30 | 34.3 | 2.7 | 19 | 40.6 | 2.3 | 42 | 48.6 | 2.1 | 39 | 95:5/65:35=0.0015 |

**Supplementary table 5.** Statistics of **Figure 3g, j** and k and **Supplementary figure 3b, h-k**. All data represent means ± SEM. * p < 0.05; ** p < 0.01; *** p < 0.001. Paired T-test or Wilcoxon matched-pairs signed rank test was performed between network activity pre, and post treatment. P-values were corrected for multiple comparisons using Bonferroni’s. Other comparisons were non-significant. NBD= Network burst duration, MFR= Mean firing rate, n= number of wells, DIV= Days in vitro.

|  |  |  | Pre-treatment |  | Post treatment |  | n | p-value |
| --- | --- | --- | --- | --- | --- | --- | --- | --- |
|  |  |  | Mean | SEM | Mean | SEM |  |  |
| **Fig. 3g** | NBD (ms) | DIV 49 | 1.01 | 0.05 | 1.33 | 0.08 | 15 | 0.0017 |
| **Fig. 3j** | NBD (ms) | 75:25 | 897.2 | 26.21 | 1108 | 37.94 | 18 | <0.0001 |
| **Fig. 3k** | NBD (ms) | 65:35 | 727.2 | 41.23 | 967.7 | 76.26 | 15 | 0.001 |
| **Sup. Fig. 3b** | MFR (Hz) | DIV49 | 12.32 | 2.48 | 2.81 | 0.58 | 8 | 0.0021 |
|  | NBR (b/min) | DIV49 | 4.05 | 0.72 | 0 | 0 | 8 | 0.0078 |
| **Sup. Fig. 3h** | MFR (Hz) | DIV 49 | 1.12 | 0.04 | 1.49 | 0.10 | 15 | 0.0044 |
| **Sup. Fig. 3i** | MFR (Hz) | 65:35 | 3.28 | 1.37 | 4.78 | 2.18 | 15 | 0.0003 |
| **Sup. Fig. 3j** | Norm. MFR | ChR2 in Ngn2 | 1 | - | 3.87 | 1.66 | 8 | 0.0391 |
| **Sup. Fig. 3k** | Norm. MFR | ChR2 in Ascl1 | 1 | - | 0.81 | 0.13 | 7 | 0.0319 |

**Supplementary table 6.** Statistics from **Figure 4**. All data represent means ± SEM. * p < 0.05; ** p < 0.01; *** p < 0.001. GABAergic sPSC amplitude and frequency were compared using non-parametric Mann-Whitney U ranked sum test. MEA parameters were compared using Mann-Whitney test with post hoc Bonferroni correction for multiple testing. NBD= Network burst duration, MFR= Mean firing rate, PRS= percentage of random spikes, n= number of wells, DIV= Days in vitro.

|  | DIV | Control |  |  | CDH13-knockdown |  |  | p-value |
| --- | --- | --- | --- | --- | --- | --- | --- | --- |
|  |  | Mean | SEM | n | Mean | SEM | n |  |
| **GABAergic SPSC amplitude** | 70 | 12.76 | 2.29 | 7 | 17.43 | 4.05 | 5 | P<0.05 |
| **GABAergic SPSC frequency** | 70 | 0.64 | 0.31 | 7 | 2.14 | 1.90 | 5 | n.s. |
| **NBD (ms)** | 49 | 751.16 | 17.53 | 39 | 613.40 | 25.37 | 24 | <0.0001 |
| **MFR (Hz)** | 49 | 3.68 | 0.31 | 38 | 2.66 | 0.36 | 33 | 0.001 |
| **PRS (%)** | 49 | 47.27 | 2.32 | 39 | 67.74 | 2.69 | 33 | <0.0001 |

**Supplementary table 7.** Statistics of **Figure 5h-n.** All data represent means ± SEM. * p < 0.05; ** p < 0.01; *** p < 0.001. Depending on normal distribution, either an unpaired T-test or Mann Whitney-U test is performed between conditions. DIV= Days in vitro.

| Panel |  | T60/T0 | Compared to | T60/T0 | p-value |
| --- | --- | --- | --- | --- | --- |
| **G** | mCherry | 0.94 ± 0.01 | CHD2 | 0.58 ± 0.01 | 0.0022 |
| **G** | mCherry | 0.94 ± 0.01 | CDH13 | 0.68 ± 0.01 | 0.0004 |
| **G** | CDH2 | 0.58 ± 0.01 | CDH13 | 0.68 ± 0.01 | 0.0001 |
| **H** | CDH13 | 0.68 ± 0.01 | GABAAα1/β3 | 0.79 ± 0.04 | 0.0077 |
| **H** | CDH13 | 0.68 ± 0.01 | GABAAα1/β3+<br>CDH13 | 0.78 ± 0.06 | 0.0253 |
| **I** | CDH13 | 0.68 ± 0.01 | ITGβ1 | | 0.0256 |
| **I** | CDH13 | 0.68 ± 0.01 | ITGβ1+ CDH13 | 0.62 ± 0.02 | 0.0350 |
| **I** | ITGβ1 | | ITGβ1+ CDH13 | 0.62 ± 0.02 | 0.0476 |
| **J** | CDH13 | 0.68 ± 0.01 | ITGβ3 | | <0.0001 |
| **J** | CDH13 | 0.68 ± 0.01 | ITGβ3+ CDH13 | | 0.0298 |
| **J** | ITGβ3 | | ITGβ1+ CDH13 | | 0.0008 |
| **L** | ITGβ3 | | GABAAα1/β3 +<br>ITGβ3 | | 0.0466 |
| **M** | CDH13+ ITGβ1/<br>GABAAα1/β3 | 0.63 ± 0.004 | mCherry | 0.94 ± 0.01 | 0.0238 |
| **M** | CDH13+ ITGβ1/<br>GABAAα1/β3 | 0.63 ± 0.004 | CDH13+ITGβ3/<br>GABAAα1/β3 | 0.93 ± 0.06 | 0.0065 |

**Supplementary table 8.**
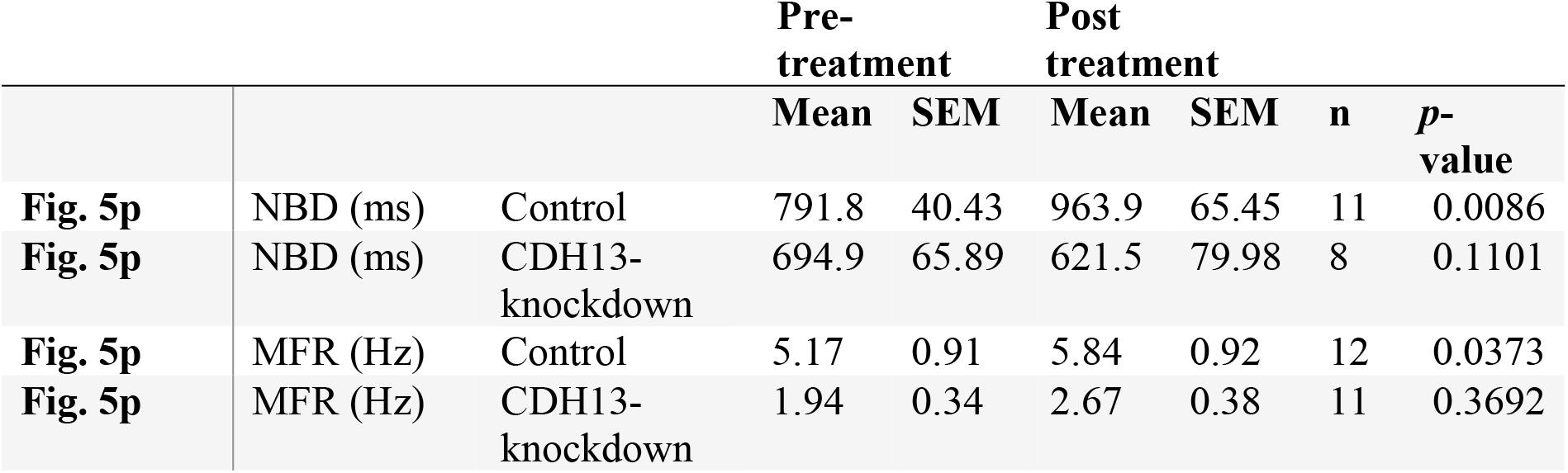
Statistics of **Figure 5o, p.** All data represent means ± SEM. * p < 0.05; ** p < 0.01; *** p < 0.001. Paired T-test was performed between pre and post-treatment conditions at DIV 49. DIV= Days in vitro.

**Supplementary table 9.**
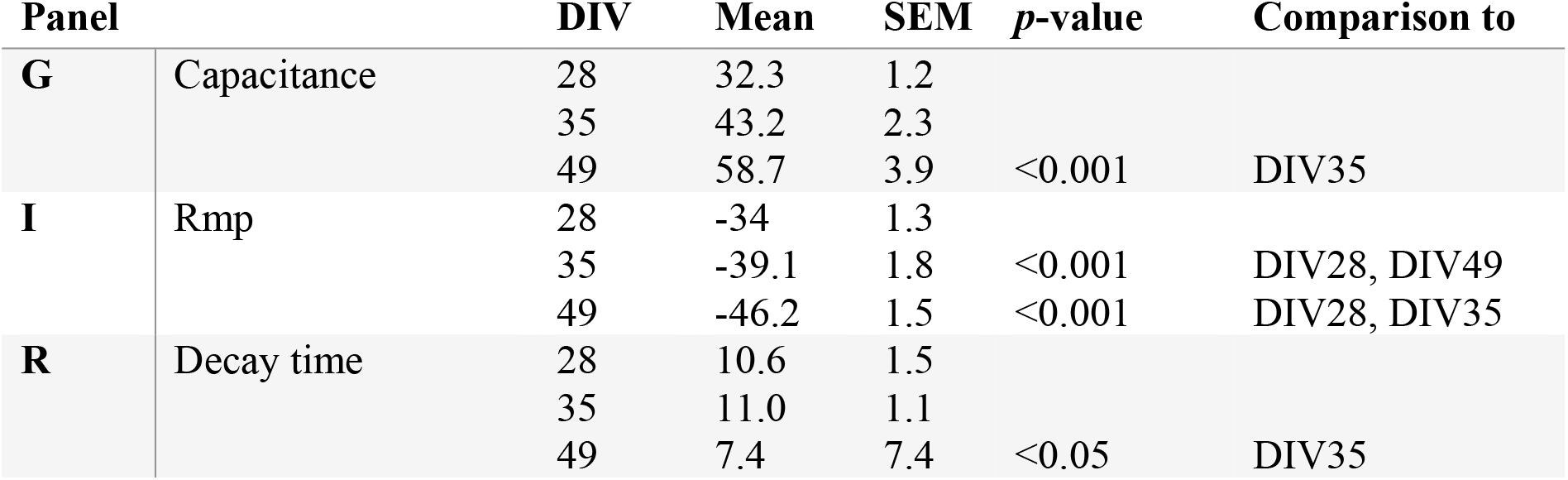
Statistics on intrinsic properties from **iGLU_Ngn2_ neurons in supplementary figure 1**. Sample size for DIV 28 n=42, DIV 35 n=40, DIV 49 n=44 cells from 3 batches. All data represent means ± SEM. * p < 0.05; ** p < 0.01; *** p < 0.001. Two-way ANOVA with Tukey correction for multiple testing was used to compare between DIVs. DIV= Days in vitro.

**Supplementary table 10.** Statistics of **supplementary figure 4c**. All data represent means ± SEM. Mann-Whitney test was performed between control and CDH13-knockdown networks. Significance was corrected for multiple comparisons using Bonferroni’s.

|  | Control |  | CDH13-knockdown |  |  |  |
| --- | --- | --- | --- | --- | --- | --- |
|  | Mean | SEM | Mean | SEM | n | *p*-value |
| **CDH13 mRNA expression** | 1 | - | 0.37 | 0.13 | 11 | 0.0091 |

**Supplementary table 11.**
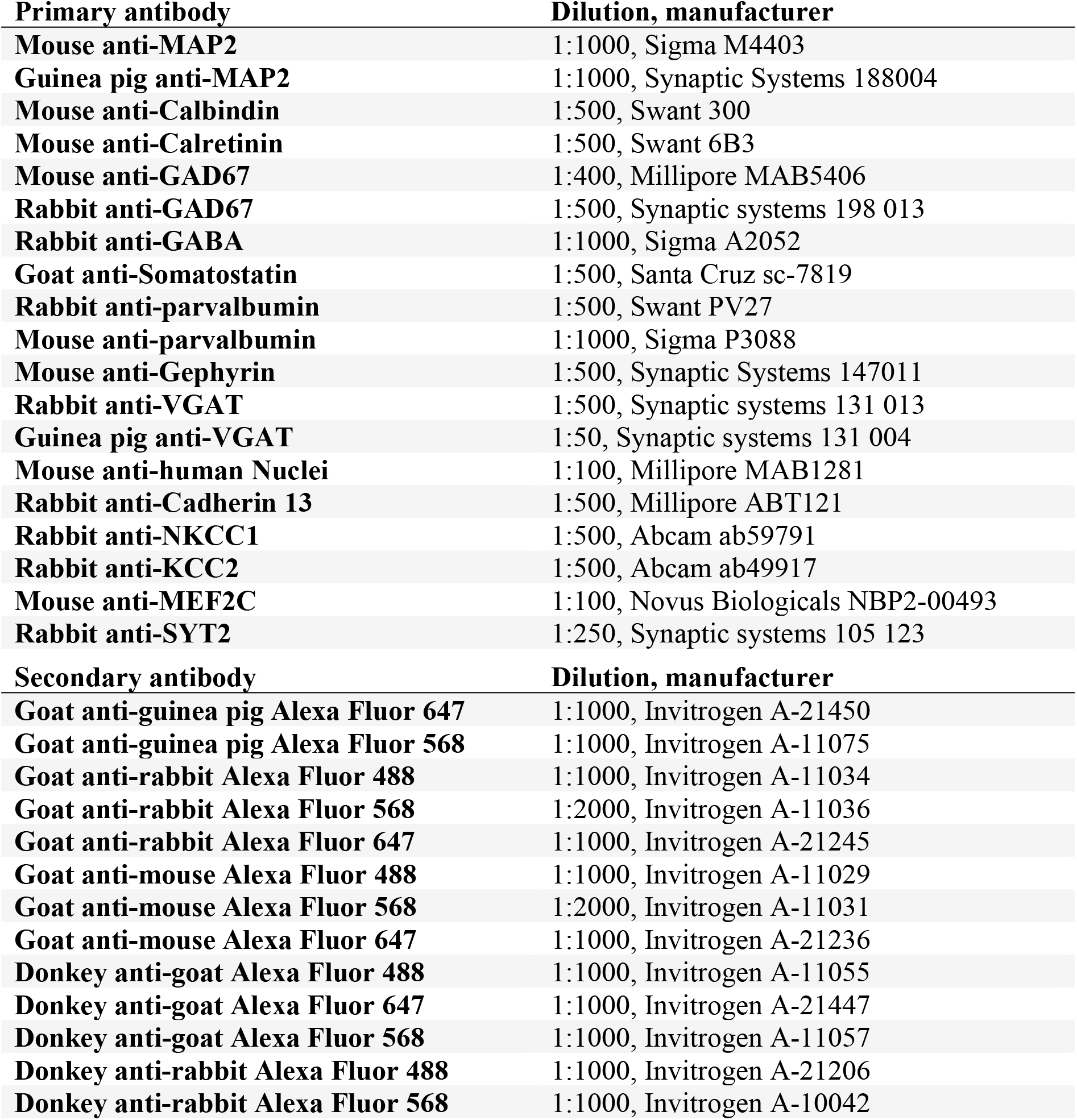
Used primary and secondary antibodies

| **Primary antibody** | **Dilution, manufacturer** |
| --- | --- |
| **Mouse anti-MAP2** | 1:1000, Sigma M4403 |
| **Guinea pig anti-MAP2** | 1:1000, Synaptic Systems 188004 |
| **Mouse anti-Calbindin** | 1:500, Swant 300 |
| **Mouse anti-Calretinin** | 1:500, Swant 6B3 |
| **Mouse anti-GAD67** | 1:400, Millipore MAB5406 |
| **Rabbit anti-GAD67** | 1:500, Synaptic systems 198 013 |
| **Rabbit anti-GABA** | 1:1000, Sigma A2052 |
| **Goat anti-Somatostatin** | 1:500, Santa Cruz sc-7819 |
| **Rabbit anti-parvalbumin** | 1:500, Swant PV27 |
| **Mouse anti-parvalbumin** | 1:1000, Sigma P3088 |
| **Mouse anti-Gephyrin** | 1:500, Synaptic Systems 147011 |
| **Rabbit anti-VGAT** | 1:500, Synaptic systems 131 013 |
| **Guinea pig anti-VGAT** | 1:50, Synaptic systems 131 004 |
| **Mouse anti-human Nuclei** | 1:100, Millipore MAB1281 |
| **Rabbit anti-Cadherin 13** | 1:500, Millipore ABT121 |
| **Rabbit anti-NKCC1** | 1:500, Abcam ab59791 |
| **Rabbit anti-KCC2** | 1:500, Abcam ab49917 |
| **Mouse anti-MEF2C** | 1:100, Novus Biologicals NBP2-00493 |
| **Rabbit anti-SYT2** | 1:250, Synaptic systems 105 123 |
| **Secondary antibody** | **Dilution, manufacturer** |
| **Goat anti-guinea pig Alexa Fluor 647** | 1:1000, Invitrogen A-21450 |
| **Goat anti-guinea pig Alexa Fluor 568** | 1:1000, Invitrogen A-11075 |
| **Goat anti-rabbit Alexa Fluor 488** | 1:1000, Invitrogen A-11034 |
| **Goat anti-rabbit Alexa Fluor 568** | 1:2000, Invitrogen A-11036 |
| **Goat anti-rabbit Alexa Fluor 647** | 1:1000, Invitrogen A-21245 |
| **Goat anti-mouse Alexa Fluor 488** | 1:1000, Invitrogen A-11029 |
| **Goat anti-mouse Alexa Fluor 568** | 1:2000, Invitrogen A-11031 |
| **Goat anti-mouse Alexa Fluor 647** | 1:1000, Invitrogen A-21236 |
| **Donkey anti-goat Alexa Fluor 488** | 1:1000, Invitrogen A-11055 |
| **Donkey anti-goat Alexa Fluor 647** | 1:1000, Invitrogen A-21447 |
| **Donkey anti-goat Alexa Fluor 568** | 1:1000, Invitrogen A-11057 |
| **Donkey anti-rabbit Alexa Fluor 488** | 1:1000, Invitrogen A-21206 |
| **Donkey anti-rabbit Alexa Fluor 568** | 1:1000, Invitrogen A-10042 |

**Supplementary table 12.**
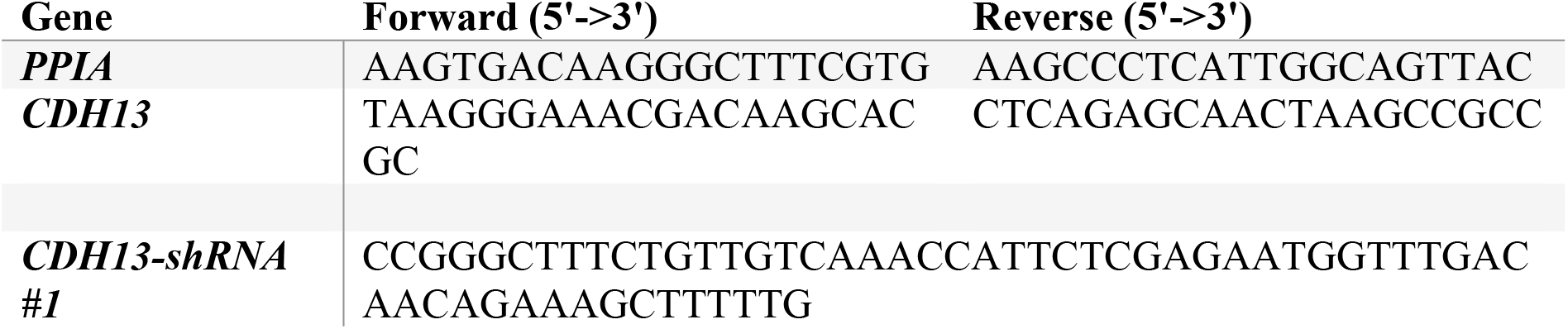
Forward and reverse primer sequences and CDH13 targeting short hairpin RNA sequences.

